# Synthetic Combinatorial Minimisation of Cell Cycle Control in Yeast

**DOI:** 10.64898/2026.07.24.740655

**Authors:** Anastasiya Malyshava, Klaudia Ciurkot, Lucas Cañizares Alonso, Armand Grollemund, Willam M. Shaw, Matteo Barberis, Tom Ellis

## Abstract

The eukaryotic cell cycle, with its inherent regulatory redundancy, provides an ideal target for exploring genome modularisation and minimisation through synthetic genomics. Building upon principles established by the Synthetic Yeast Genome (Sc2.0) project, we used CRISPR-mediated genome engineering to relocate nine key cell cycle genes into a synthetic gene cluster in the *S. cerevisiae* genome, to allow combinatorial study of these genes in yeast. We employed Cre/loxP recombination to rapidly generate hundreds of strains with different gene deletion combinations in the module, with the objective of identifying minimal gene sets that permit robust cell cycle function. Using FACS-sorting and POLAR (Pool of Long Amplified Reads) sequencing, we conducted high-throughput analysis of gene deletion combinations in large cell pools, detecting approximately 80% of theoretically possible gene combinations, including those predicted from prior mathematical modelling studies. Our findings demonstrate that the cell cycle gene set can be minimised while maintaining viability, though only select combinations of gene deletions ensure robust fitness across different conditions. This work establishes a framework for genome minimisation, opening the door to the design of simplified, modular synthetic genomes for diverse applications.

## Introduction

Understanding the logic of life can be accelerated by studying minimal cells containing only the genes required for essential cellular functions. Advances in whole-genome engineering are now making the construction of such minimal genomes increasingly feasible^1^. While synthetic genomics initially focused on recreating and minimising bacterial genomes^2,3^, the impending completion of the Sc2.0 synthetic yeast genome^4–7^ paves the way for more radical synthetic genome redesigns, in eukaryotes like *Saccharomyces cerevisiae*. As a model organism, budding yeast offers unique advantages, combining genetic tractability and a small genome with highly conserved cell cycle mechanisms that parallel those in other eukaryotes, including humans^8,9^. These experimental advantages make yeast an ideal platform for addressing synthetic genome design challenges, particularly understanding how essential genes can be reorganised and chromosomes minimised while maintaining cellular function. Building on the success of Sc2.0, a proposed Sc3.0 project aims to create a minimal, modular synthetic yeast genome for laboratory and bioreactor environments^10^.

The eukaryotic cell cycle, a well-understood fundamental process, presents an ideal target for modularisation through synthetic genomics. In budding yeast, the cell cycle is driven by waves of cyclin/Cdk1 complexes **(Fig. 1A)**, where Cdk1 serves as the primary cell cycle regulator^11^. Nine distinct cyclins (Cln1-3 and Clb1-6) sequentially bind and activate Cdk1, conferring substrate specificity for phosphorylating diverse targets from inhibitors to transcription factors^12^. These cyclins oscillate through the cell cycle in a predictable pattern, with their sequential waves and stepwise Cdk1 activation orchestrating unidirectional cell cycle progression through distinct phases. The waves of cyclins drive essential events including S phase entry and metaphase progression, while cyclin inactivation enables cytokinesis, spindle breakdown, and licensing of replication origins for the next cell cycle^13^. Despite the presence of nine cyclin genes, significant redundancy suggests fewer, potentially even a single cyclin, may suffice for robust cell cycle progression^14–18^. The minimal number of cyclins required remains unknown and depends on various regulators.

**Figure 1:**
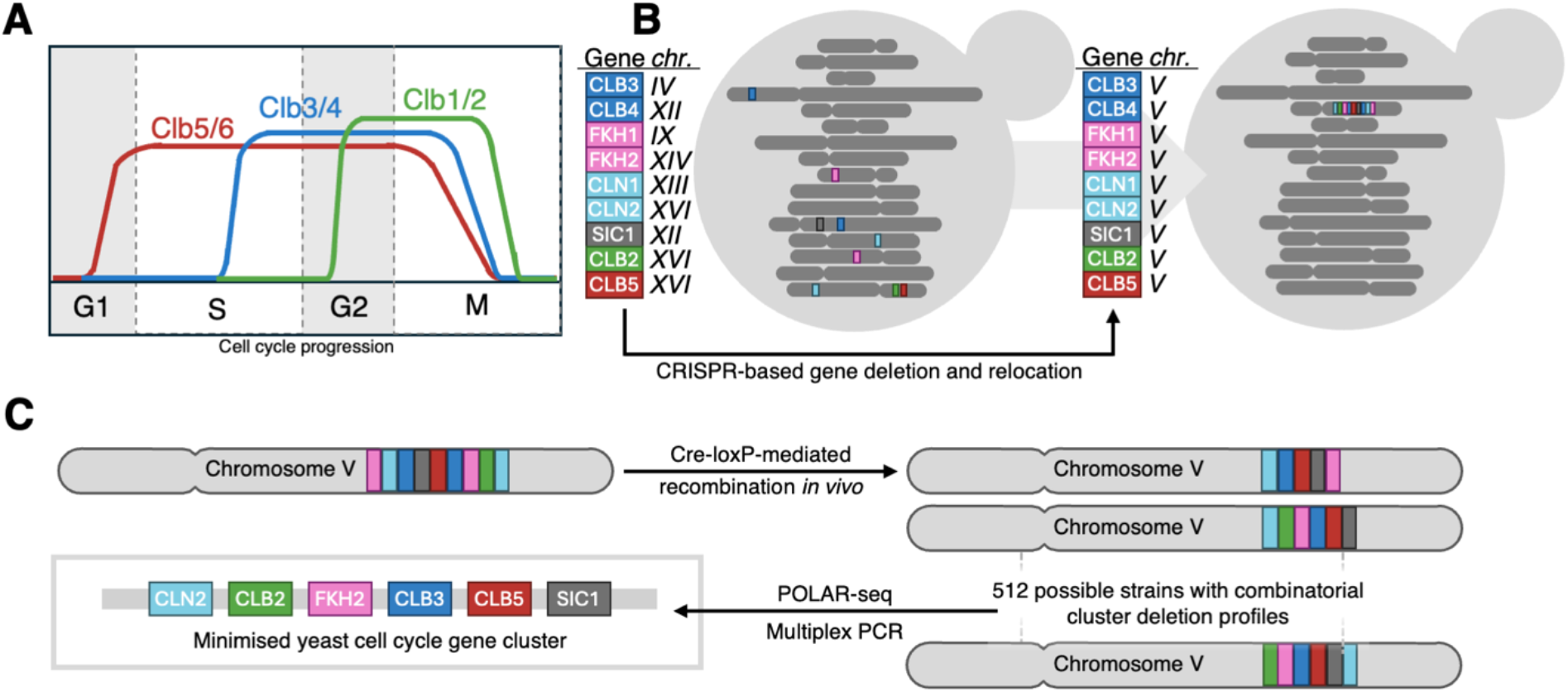
Overview of experimental strategy for determining and validating minimal cell cycle networks through synthetic genomics and systems design methodologies. (**A**) Oscillatory pattern of mitotic cyclin/Cdk1 complex activity during cell cycle progression, showing sequential waves of Clb5/6 (red colour), Clb3/4 (blue colour), and Clb1/2 (green colour). **(B)** A schematic of the *S. cerevisiae* genome shows relative chromosomal positions of selected cell cycle genes highlighted as vertical bars. CRISPR-based gene deletion and relocation of all nine genes into a single chromosomal locus was used to construct a synthetic genome module (shown as coloured vertical bars). **(C)** Generation of strains with combinatorial cluster deletions via Cre/loxP recombination, with examples of resulting genotypes where colours correspond to individual genes.

The cell cycle control system functions as a bistable, ultrasensitive biochemical switch incorporating positive and negative feedback mechanisms. Extensive studies on the budding yeast cell cycle have led to various *in silico* models simulating cell cycle dynamics and phase transitions^19–22^. These computational approaches range from complex models capturing detailed molecular interactions and stochastic fluctuations, to simplified minimal systems exploring essential regulators. Notably, computational models by Barberis’s group qualitatively reproduce B-type cyclin oscillatory behaviour using minimal gene sets^23–25^. They propose a minimal autonomous oscillator, coordinated through cyclin/Forkhead interactions, that rationalises a quantitative model of Cdk1 control in yeast, identifying a small set of regulators sufficient to reproduce the oscillatory cyclin waves^26,27^. These predictions provide us with a testable set of genes that encode a core cell function, an ideal set to consider as a testbed for minimising content in genomes and moving genes from their native loci into synthetic modules defined by function.

Here, we present a unique synthetic genomics approach to experimentally test and validate predictions of the minimal cell cycle networks that uses the principle of genome ‘defragmentation’, where genes encoding a common function are relocated from around the genome into a single module^1,28,29^ Using CRISPR-based gene deletion combined modular DNA assembly, we constructed a synthetic cell cycle control gene cluster **(Fig. 1B)**. Then, through Cre/loxP-mediated recombination, we generated diverse strains by combinatorial deletion in this synthetic cluster, analysing this using Pool of Long Amplified Reads sequencing (POLAR-seq^30^) and multiplex PCR **(Fig. 1C)**. By comparing our high-throughput experimental findings with computational models, we bridge theoretical predictions and empirical results, advancing our understanding of minimal gene requirements for cell cycle function. This knowledge represents a step towards genome designers being able to predict and assemble optimal gene sets for specific cellular functions, integrating synthetic biology and systems modelling approaches to facilitate the design of synthetic modular genomes.

## Results and Discussion

### Building Synthetic Cell Cycle Gene Clusters

The action of Cdk1 in running the cell cycle is controlled by 8 cyclins and potentially dozens of regulatory proteins. In the minimal networks proposed and simulated by Barberis, the number of genes required to achieve robust cell cycle oscillations can be reduced to just three key regulators, *FKH1*, *FKH2*, and *SIC1* regulating a subset of the 8 cyclin genes by controlling their expression (*FKH1 FKH2*) and degradation (*SIC1*)^23–27^. To establish a synthetic genomics approach to combinatorially explore the minimal set of genes controlling cyclin wave dynamics, we thus focused on relocating these three key regulators and the 8 cyclins: two G_1_ cyclins (*CLN1* & *CLN2*), the six B-type cyclins (*CLB1* to *CLB6*), from their native loci to a synthetic gene cluster, by gene deletion and gene reinsertion at the cluster locus. Based on insights from a large-scale trigenic interaction study^31^, we strategically deleted two to three native genes at a time to minimise complex genetic interactions.

For gene deletion we used modular CRISPR/Cas9 methods for yeast to systematically remove open reading frames of the target genes, replacing them with unique 24 bp barcode sequences^32,33^. We then iteratively constructed a synthetic gene cluster, integrating DNA along with auxotrophic markers (*URA3*, *LEU2*, or *HIS3*) to allow for selection of strains with integrated clusters. This process generated a series of increasingly engineered strains containing clusters of 3, 5, 6, and finally 9 cell cycle genes **(Fig. 2A)**. Given the essential roles and complex regulation of these genes, we relocated all genes with their flanking native regulatory elements (promoters, terminators) in order to try to preserve their dynamic regulation^34,35^. The fidelity of gene deletions and cluster assembly was verified through colony PCR and sequencing. While we initially targeted all cyclin genes, technical difficulties in deleting *CLB1* and *CLB6* (a gene pair naturally co-located at a single locus) led us to proceed with minimisation experiments using the 9-gene cluster (9KO9KI) as our base strain. For more details of the strain construction approach taken, see **Fig. S1, Fig. S2** and the **Supplementary Note**.

**Figure 2:**
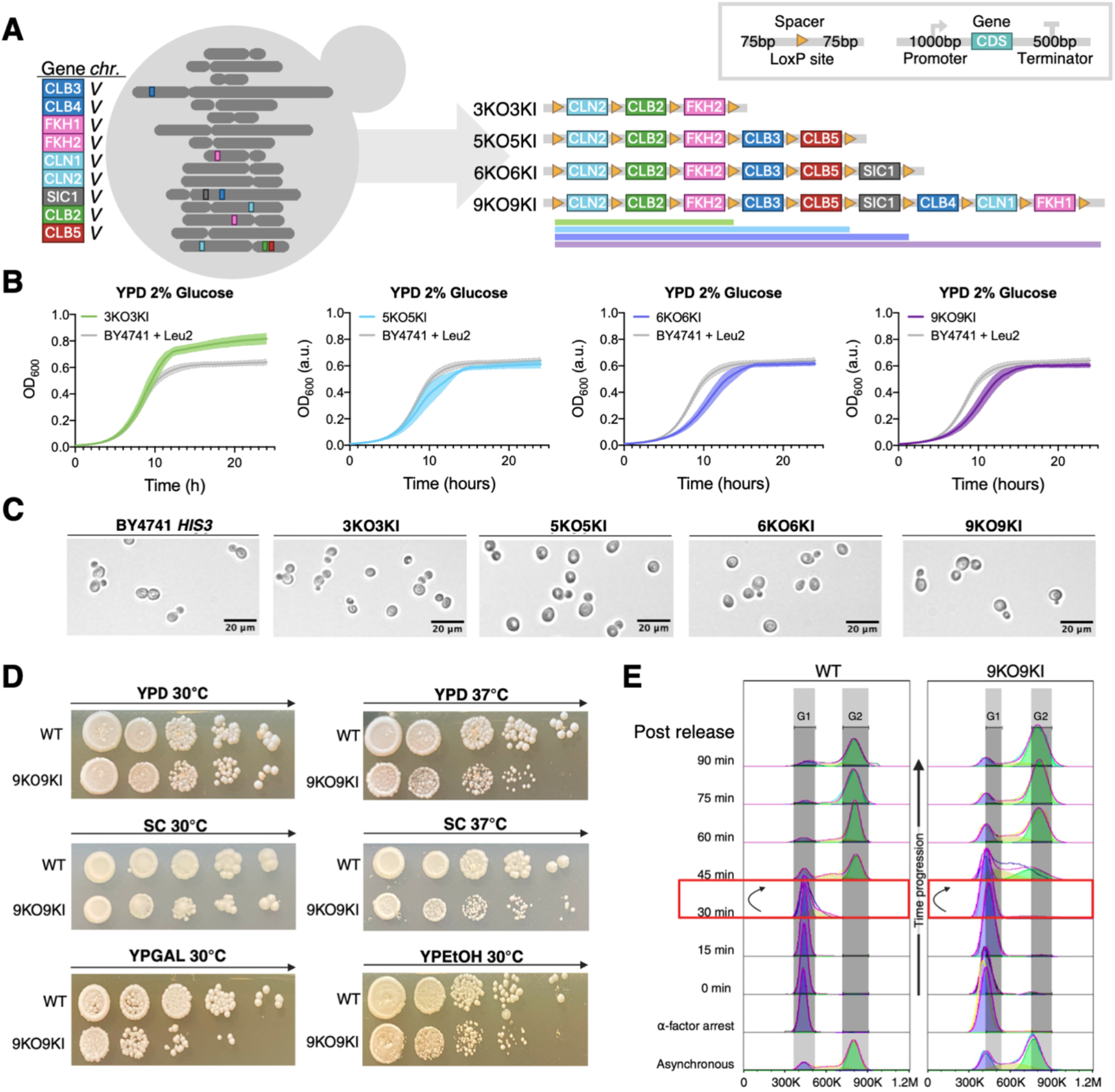
Construction and characterisation of synthetic cell cycle gene clusters in *S. cerevisiae*. (**A**) Schematic of CRISPR/Cas9-mediated construction of synthetic cell cycle gene clusters. Stepwise gene deletion and cluster assembly is indicated by colours: green for 3-gene cluster (3KO3KI: *CLN2*, *CLB2*, *FKH2*), blue for 5-gene cluster (5KO5KI: adding *CLB3*, *CLB5*), lavender for 6-gene cluster (6KO6KI: adding *SIC1*), and purple for 9-gene cluster (9KO9KI: adding *CLB4*, *CLN1*, *FKH1*). Inset: structure of each integrated gene with native regulatory elements and the spacers with loxP sites flanking the genes in all clusters. **(B)** Growth curves of the wild-type (WT) and cluster strains in YPD medium containing 2% glucose at 30°C in 96-well plates. Mean OD_600_ values from 5 biological replicates are plotted as lines; the error bars represent SEM and are drawn as dotted lines with semi-transparent fill areas. The BY4741 strain with *LEU2* reintegrated at the *URA3* locus (strain yAM075) was used as a control. OD_600_ measured every 15 min using a microplate reader. **(C)** Representative bright-field microscopy images of WT and synthetic cluster strains at 20x magnification (scale bar = 20 μm) grown in YPD 2% glucose media. **(D)** Spot assay viability assessment of WT and 9KO9KI strains. Ten-fold serial dilutions of normalised cultures (OD_600_ = 1.0) were spotted from left to right on media with different carbon sources: 2% glucose (YPD and SC media), 2% ethanol (YPEtOH), and 2% galactose (YPGAL) and incubated at 30°C or 37°C for 3 days. n=3 for the 9-gene cluster and the control strain yAM075 (BY4741 with *LEU2* reintegrated at the *URA3* locus); the representative images shown were cropped from the same plates (full data in **Fig. S3**). **(E)** Flow cytometry analysis of cell cycle progression in *bar1* knocked-out WT and 9KO9KI strains following α-factor synchronisation and release (0-90 min). Representative flow cytometric histograms overlaid half-offset for each time point are shown (n=2 replicates); cell counts (y-axis) are plotted against DNA content (x-axis, BL1-H). Asynchronous cultures were used as controls for constraining G_1_ and G_2_ (shown as grey vertical boxes) and the Watson Pragmatic model for cell cycle analysis was applied to fit a curve to the SYTOX Green fluorescence data. G_1_ phase, S phase, and G_2_/M phase are denoted by blue, yellow, and green, respectively. Red box marks the G_1_-to-S transition time point, with arrows comparing WT and 9KO9KI strains, revealing extended G_1_ in 9KO9KI.

### Phenotypic Characterisation of the Yeast Synthetic Cell Cycle Cluster Strains

We observed a gradual reduction in growth rate with each cluster expansion round **(Fig. 2B, 2C Fig. S3)**. While the 3-gene and 5-gene cluster strains grew similarly to wild type, the 6-gene and 9-gene strains showed slightly delayed entry into logarithmic growth in 2% glucose media **(Fig. 2B)**. This growth defect was further pronounced in non-glucose carbon sources, particularly in galactose and ethanol media **(Fig. S3)**. Microscopy revealed slightly larger cells in the 5-, 6– and 9-gene cluster strains compared to wild type and the 3-gene cluster **(Fig. 2C, Fig. S3)**, with spot assays indicating lower colony count for the 9-gene cluster in low and intermediate OD_600_ dilutions under various conditions **(Fig. 2D, Fig. S3)**.

To assess cell cycle progression, we synchronised cells with α-factor after deleting *BAR1* in our cluster strains^36^. Flow cytometry cell cycle analysis revealed an extended G_1_ phase in both the 6-gene and 9-gene cluster strains **(Fig. 2E, Fig. S4)**, lasting up to 60 and 45 minutes, respectively. Whole genome sequencing analysis, conducted to investigate potential off-target mutations, revealed unexpected aneuploidisation in the 9-gene cluster strain **(Fig. S5, Fig. S6)**. Aneuploidy in yeast is a known cause for G1 phase delay in the cell cycle^37,38^ and while yeast can undergo ploidy alterations as a consequence of stress^39–41^, it may simply be that deletion of key genes (e.g. *CLB2* and *SIC1*) in steps during synthetic cluster construction disrupted faithful chromosome segregation in intermediate strains, causing all downstream strains to inherit aneuploidy^42,43^.

### Combinatorial Minimisation of the Yeast Cell Cycle Gene Cluster

To investigate minimisation outcomes in our synthetic cell cycle gene cluster, we adapted the SCRaMbLE (Synthetic Chromosome Rearrangement and Modification by loxP-mediated Evolution) method from the Sc2.0 project^44^. Our gene module comprised cyclins and regulators flanked by loxP sites, enabling estradiol-induced Cre-EBD enzyme to randomly delete DNA between these sites **(Fig. 3A)**. While the system predominantly generated deletions between loxP sites in cis arrangement, rare duplications (<0.004%) could arise through trans recombination between sister chromatids post-replication. With nine genes that could be either present or absent, this design enabled 2^9^ (512) possible combinations, though some would be lethal given the module’s overall essentiality — providing an ideal system for studying viable deletion combinations.

**Figure 3:**
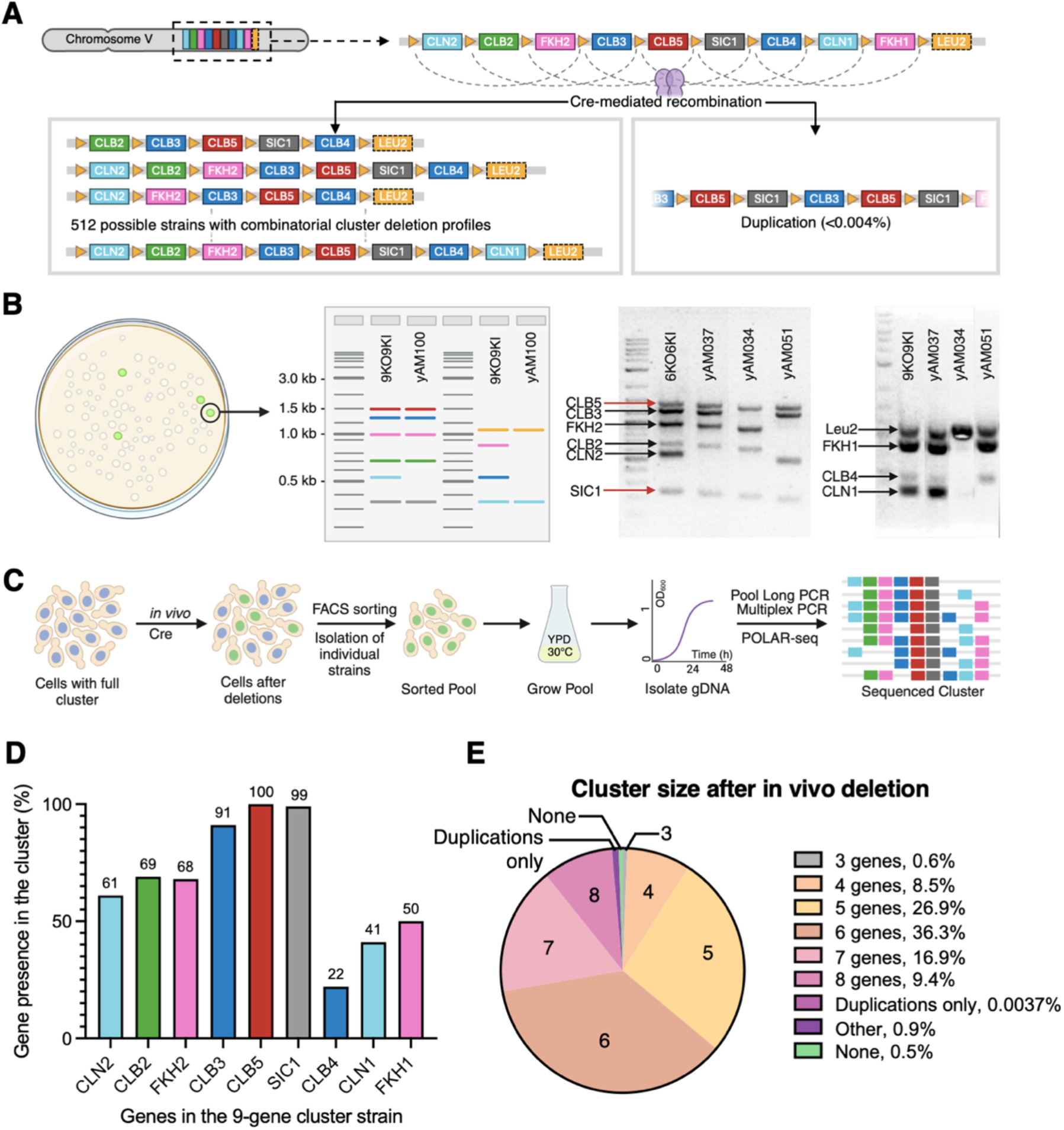
Combinatorial minimisation of a synthetic yeast cell cycle gene cluster *in situ*. (**A**) Nine cell cycle genes flanked by loxP sites (orange triangles) undergo Cre-mediated recombination. Cre dimers (light blue) trigger deletions (cis) or rare duplications (trans, <0.004%) between loxP sites facing the same direction, enabling 512 possible combinations. **(B)** Genotyping of minimised strains using Multiplex PCR. Fluorescent colonies are screened using two PCR reactions: one detecting six genes (*CLB5*, *CLB3*, *FKH2*, *CLB2*, *CLN2*, *SIC1*) and another for three genes plus marker (*LEU2*, *FKH1*, *CLB4*, *CLN1*). Representative gel images from strains yAM037, yAM034, and yAM051 with 1, 5, and 3 gene deletions, respectively, show consistent retention of *CLB5* and *SIC1*. **(C)** High-throughput workflow combining FACS sorting of minimised strains with POLAR-seq analysis. FACS-sorted GFP-positive cells are grown (20°C, 48h; then 30°C, 48h) before pooled gDNA isolation. High molecular weight gDNA templates are used for long-range PCR amplification of the cluster region, followed by Oxford Nanopore sequencing of the amplicon library. **(D)** Mean gene retention frequencies in minimised clusters from sequencing data, showing preferential retention of *CLB5* (100%) and *SIC1* (99%) based on gene occurrence count in all generated reads. **(E)** Distribution of cluster sizes after *in vivo* deletion, determined by MinION sequencing.

We initially screened recombinant genotypes using a Cre-dependent GFP reporter assay as an indirect indicator of genomic rearrangements^29^. DNA from green fluorescent colonies was extracted for Multiplex PCR analysis **(Fig. 3B)**, which employed a loxPsite targeting forward primer and multiple reverse primers to generate a diagnostic ladder-like pattern on gel electrophoresis, where missing bands indicated gene deletions **(Fig. 3B)**. The reaction was split into two: one for six genes (*CLB5*, *CLB3*, *FKH2*, *CLB2*, *CLN2*, *SIC1*) and another for the remaining genes (*CLB4*, *CLN1*, *FKH1*) plus *LEU2* marker. While analysis of 92 colonies provided initial insights into rare deletions of *CLB5* and *SIC1* **(Fig. 3B, Fig. S7)**, this method proved labour-intensive for comprehensive library screening.

To achieve higher throughput, we used FACS to isolate >3×10^5^ recombined cells using the same fluorescent reporter, despite low GFP-positive cell numbers that likely reflected synthetic lethality among cell cycle gene deletions **(Fig. S8)**. From this diverse pool of recombinants, we extracted high-quality genomic DNA for subsequent analysis using POLAR-seq **(Fig. 3C)**, a high-throughput method that combines long-range PCR amplification of the synthetic cluster region (up to ∼28 kb) with long read nanopore DNA sequencing^30^. This approach captured reads spanning the entire cluster, revealing deletion combinations occurring *in vivo* at single-cell resolution. Multiplex PCR on FACS-sorted colonies further validated these findings. This integrated approach allowed us to rapidly and cost-effectively sequence large libraries of recombined cluster strains, detecting ∼80% of all theoretically possible combinations, including rare variants that would be difficult to identify otherwise.

### Frequencies of Individual Gene Deletions in the Cell Cycle Gene Cluster

POLAR-seq analysis revealed that every gene was deleted in at least one strain. While *CLB4*, *CLN1*, and *FKH1* most frequently deleted **(Fig. 3D)**, *CLB5* and *SIC1* showed remarkably high retention rates (∼100% and 98.55% respectively), reflecting their crucial role in the G_1_/S transition^45^. This preferential retention proved especially interesting as these genes occupy central positions in the cluster were, theoretically, higher deletion frequencies would be expected due to proximity to multiple loxP sites. Their persistence suggests strong selective pressure for maintenance, though distinguishing between positional effects and functional essentiality would require further experiments involving repositioning these genes within the cluster. The reduced expression of *SIC1* in our synthetic cluster may also contribute to this retention pattern **(Fig. S9)**.

Deletion frequencies among paralogs correlated with their relative protein abundances^46,47^, showing higher retention of *CLN2* over *CLN1* (61% vs. 41%) and *CLB3* over *CLB4* (91% vs. 22%). *FKH1* and *FKH2* showed similar deletion frequencies, possibly due to their indirect role as transcription factors^48–50^. These findings largely aligned with the ‘Barberis model’ predicting *CLN2*, *CLB5*, *CLB3*, *CLB2*, *SIC1*, and *FKH2* as the minimal gene set for autonomous cell cycle oscillations, although the absence of *CLB1* and *CLB6* from our synthetic cluster limits conclusions on cyclin pair redundancy.

We identified 432 distinct genotypes through long-read sequencing, capturing ∼84% of theoretically possible combinations and likely representing most viable minimised clusters after accounting for synthetic lethal interactions (see **Table S1** for full list of genotypes). Most viable strains retained 5-8 genes, with three– and four-gene deletions being most frequent (36% and 27%, respectively) **(Fig. 3E)**. The low frequency of parental 9-gene clusters (0.51% of sequenced amplicons) confirmed strong correlation between GFP reporter activation and cluster recombination. Of the identified genotypes, 4.4% showed both deletions and duplications.

Larger clusters were rare: 9-gene variants comprised 0.86% of sequenced amplicons (0.51% parental, 0.35% with combined deletions and duplications), while expanded clusters with 10-11 genes represented <0.5% of reads and those with 12-13 genes constituted <0.004% of reads. This low frequency of duplications confirms the system’s strong bias toward deletions, indicating that duplications did not significantly impact our analysis of deletion profiles.

### Genotypes of Combinatorically Rearranged Synthetic Clusters

We next analysed the ten most abundant genotypes found in post-Cre/loxP cell pools using the long-read sequencing data. The top 10 genotypes accounted for 87.4% of all annotated reads and all contained between one to four gene deletions **(Fig. 4A)**. The three most prevalent genotypes each represented around 15% of reads, while genotypes 4-7 and 8-10 each represented around 5% and 3% of reads respectively.

**Figure 4:**
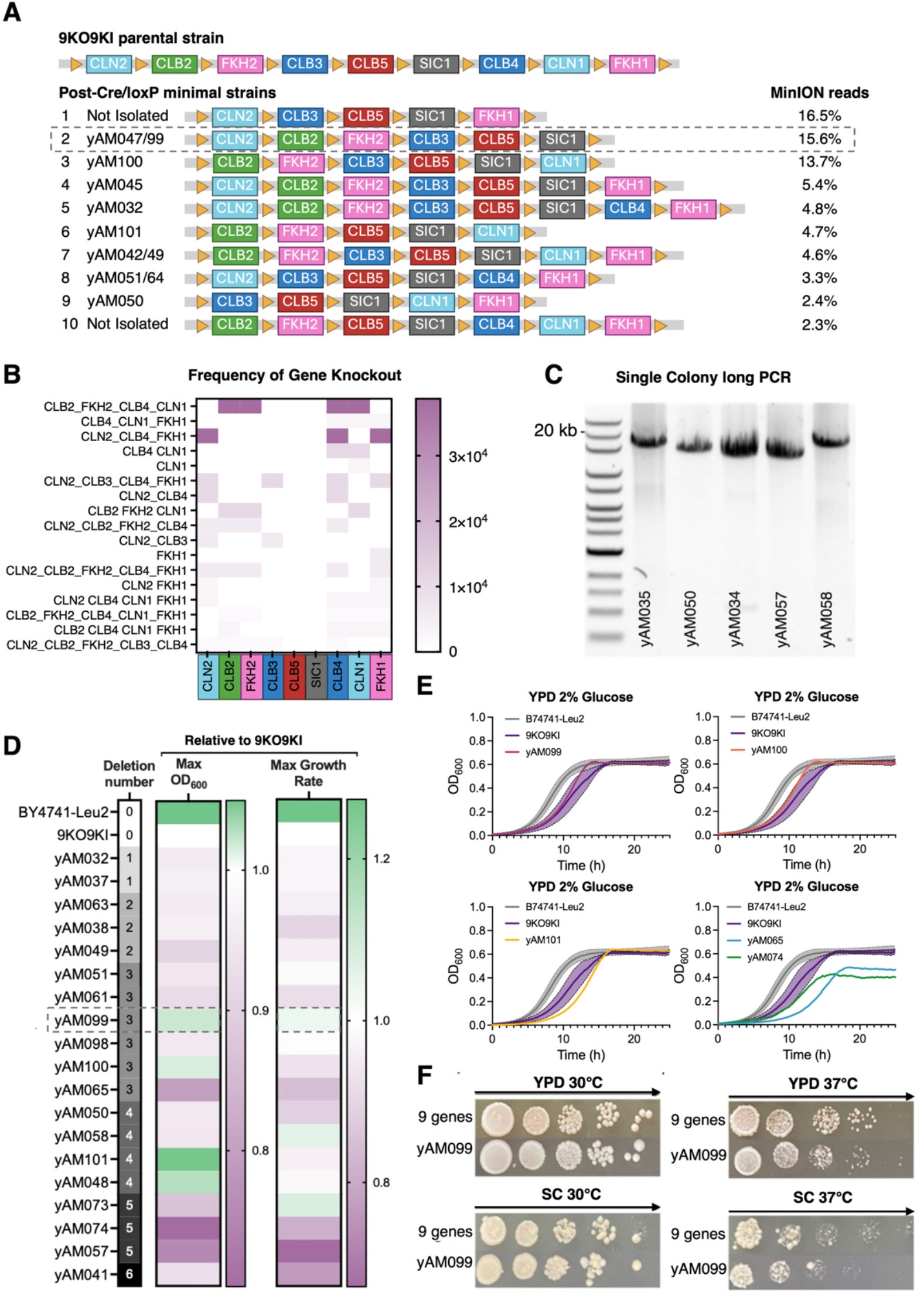
Genotypic and phenotypic characterisation of minimised cell cycle strains. (**A**) Top ten most abundant genotypes in the pool of post-Cre/loxP cells sequenced on MinION platform. Parental 9KO9KI strain shown at top; coloured boxes indicate retained genes, with corresponding independently isolated strains listed on left. Genotypes ranked by sequencing read abundance. **(B)** Heatmap showing frequency of gene deletion combinations in minimised strains, with knockout combinations (y-axis) and cluster genes (x-axis) for top 20 genotypes from long read sequencing. Combinations ordered by abundance; colour intensity represents read count. **(C)** Long PCR validation of selected minimised strains showing variable cluster sizes. 0.4% agarose gel was used with the NEB Quick-Load^®^ 1 kb Extend DNA Ladder. **(D)** Growth performance matrix of minimised strains showing maximum OD_600_ of growth curves and maximum growth rate, normalised to the 9-gene cluster (9KO9KI) strain values. Growth measured in YPD for 48 h from initial OD_600_ of ∼0.005, with measurements every 15 min (n=3). Strains ordered by number of cluster genes deleted. Colour scale: green represents faster growth rate and higher maximum OD than the 9K09KI strain, white is equivalent and purple is slower growth rate and lower maximum OD. **(E)** Growth curves in YPD 2% glucose (30°C) comparing selected minimal strains with BY4741-*LEU2* and 9KO9KI controls. OD_600_ measured every 15 min using a microplate reader. Control strains: n=5 biological replicates, mean ± SEM shown as lines with shaded areas; minimal strains: n=3 technical replicates. Values normalised to media blank. **(F)** Spot assays comparing growth of the yAM099 minimal strain with the parental 9-gene cluster strain. Ten-fold serial dilutions (starting OD_600_=1.0) on YPD and SC media with 2% glucose, incubated at 30°C and 37°C for 3 days. Representative images shown; n=3. Images cropped from the same plates.

Six of the ten most abundant genotypes retained one CLN gene, *SIC1*, and one FKH gene, while four retained both FKH genes. The top three genotypes, together constituting ∼47% of reads, closely resembled the Barberis model, containing one CLN gene, *SIC1*, one FKH gene, and at least two or three CLB genes retained from the cluster. A less frequent variant of the first genotype (yAM050, genotype 9) showed *CLN2* deletion instead of *CLN1*, consistent with our observation of *CLN1* being deleted more frequently than *CLN2* (61% vs 41% in **Fig. 3D**).

While our long-range PCR and sequencing analysis estimated combinatorial deletion genotype frequencies in the yeast cell pool, it did not directly measure strain fitness. The use of long PCR may also introduce bias in these results, as PCR preferably amplifies shorter DNA fragments, an anticipated limitation of the current POLAR-seq method^30^. Interestingly, when picking yeast from individual colonies from our library, no strains were isolated that contain the most abundant genotype from our POLAR-seq data (genotype 1, **Fig. 4A**), challenging the assumption that this genotype is the most common in the post-Cre/loxP cell population.

### Combinatorial Deletions in the Pools of Post-Cre/loxP Cells

POLAR-seq of post-Cre/loxP cell pools allowed us to explore combinatorial deletions of non-essential cell cycle genes in the 9-gene cluster strain **(Fig. 4B)**. POLAR-seq analysis revealed three predominant deletion combinations in post-Cre/loxP cell pools: Δ*cln1*, Δ*fkh2*, Δ*clb4*, and Δ*clb2* (16.5% of reads); Δ*cln1*, Δ*fkh1*, and Δ*clb4* (15.6%), and Δ*cln2*, Δ*fkh1*, and Δ*clb4* (13.7%). Notably, a variant of the most common deletion, lacking the Δ*cln2* gene (Δ*cln2*, Δ*fkh2*, Δ*clb4*, Δ*clb2*), was also identified in all sequencing datasets, comprising 2.4% of genotype reads.

Further analysis of deletion patterns identified common combinations in the sequencing dataset. The data showed frequent co-deletion of *CLN1* with *CLB4*, and *CLN2* with *CLB4* **(Fig. 4B)**. *FKH2* and *CLB2* were also frequently co-deleted. Co-deletion frequency may be directly related to the functional relationships of the genes or could be due to genes being physically in the synthetic gene cluster and thus more likely to be co-deleted by the Cre recombinase.

In contrast, some gene pairs were rarely deleted together, most notably the neighbouring genes *CLB5* and *SIC1* (<0.5-0.1% of reads), possibly due to their early cell cycle activation and critical roles in cellular fitness^45^. Interestingly, when examining single-gene deletions in the top 20 genotypes, Δ*cln1* and Δ*fkh1* knockouts were most common, an observation that, together with our earlier findings **(Fig. 3D)**, supports that cells may maintain viability with either paralog from the *CLN1/2* or *FKH1/2* pairs.

### Phenotypic Characterisation of Selected Minimised Strains

To investigate how these gene deletion combinations affect cellular fitness, we isolated and characterised minimal cell cycle strains from independent SCRaMbLE experiments. We first genotyped 92 colonies by performing Multiplex PCR analysis of lysed colonies from isolated strains **(Fig. S7)**. From that we narrowed down to 25 strains of interest, amplified the cluster DNA from these strains by long range PCR (**Fig. 4C**) and used barcoded pooled nanopore sequencing of these amplicons to confirm the cluster contents of all 25 strains **(Table S2, S3)**. In total, there were 21 unique cluster genotypes among the 25 strains, as 4 pairs of strains were present with the same cluster combination, despite being isolated individually.

Growth curve experiments in YPD 2% glucose were used to analyse growth dynamics of all the individually isolated strains. We quantified maximum OD_600_ and maximum growth rate for each strain, normalised the values to the 9-gene cluster strain and visualised these data as a heatmap **(Fig. 4D)**. As can be seen from the increased purple toward the bottom of the heatmap, strains with more deletions tend to achieve lower cell densities after a growth cycle and have a reduced maximum growth rate. Considerable variation was observed in all growth parameters, particularly among strains carrying 3 to 5 deletions, indicating that specific deletion combinations, rather than gene deletion number alone, influences growth.

Notably, strain yAM099 (carrying Δ*clb4*, Δ*cln1*, Δ*fkh1* deletions) and yAM100 (carrying Δ*clb4*, Δ*cln2*, Δ*fkh1* deletions) maintained growth characteristics comparable to the 9KO9KI control strain despite having 3 genes deleted **(Fig. 4D, 4E)**. These strains closely match the Barberis minimal cell cycle model^26,27^, with *CLB1* and *CLB6* remaining in their native genomic loci as they are not part of the synthetic cluster. The comparable performance of these strains, differing only in Δ*cln1* versus Δ*cln2*, suggests alternative configurations can achieve similar fitness levels while reducing cyclin paralog redundancy. Among the cyclin B genes, deletion of *CLB4* showed minimal impact on growth, even in combination with Δ*clb3*, as demonstrated by strains yAM098 and yAM101 **(Fig. 4E)**. However, good growth showed a dependency on *SIC1* and at least one gene from each *CLN1,2* and *FKH1,2* pair. This was particularly evident in strains yAM065 and yAM074, where co-deletion of *CLN1*, *CLN2*, and *SIC1* resulted in severe growth defects **(Fig. 4E)**.

Using spot assays, we further evaluated the deletion strains alongside the parental strain. While yAM099 grew normally at 30°C on both YPD and SC media, it exhibited growth defects at elevated temperature stress (37°C) in both media types **(Fig. 4F)**. Expanding this analysis, we evaluated strain viability by developing a standardised spot-assay **(Fig. S10)** and then tested viability in various growth conditions **(Fig. S11)**. These spot assay results were then qualitatively analysed and used to generate a visual summary heatmap of strain performance in different conditions, and overall, where purple shades indicate severely impaired growth (**Fig. S12**). Most strains grew well under standard conditions, but elevated temperatures revealed clear growth defects, with many strains struggling at 37°C, regardless of the number of genes deleted from the synthetic cluster. Strains with poor growth in YPD growth media at elevated temperatures also exhibited similar growth defects in SC medium under the same conditions **(Fig. S11)**. Growth in non-glucose carbon sources was also assessed, with growth in 2% galactose and 2% ethanol revealing strains that grow poorly in different media (**Fig. S11**), and identifying minimised strains like yAM098, yAM099 and yAM058 whose growth appears robust to such changes (**Fig. S12**)

The spot assay results show that stress conditions can reveal specific gene deletion sets that affect cell cycle dynamics that may not be detectable if assays are done under standard growth conditions. It also narrowed down the number of deletion sets that can be considered close to the fitness of the parental strain. These findings confirmed that while cell cycle control can be minimised, genetic redundancy is likely to be important for maintaining robustness under diverse conditions, including alternative carbon sources and heat stress^51^. Our results show that as more cyclin genes and regulators are removed, cell growth in diverse environments becomes increasingly compromised, aligning with existing literature on genetic redundancy in the cell cycle^12,18,52–54^.

## Discussion

Our work demonstrates the application of CRISPR-based genome engineering to ‘defragment’ the yeast genome by relocating 9 functionally related cell cycle genes into a gene cluster on chromosome V. We call this cluster, and others like it, a synthetic gene module (SGM) as it is anticipated that it can be a modular component of a future minimal synthetic yeast genome^29^. The cell cycle SGM contains cyclins and transcriptional regulators flanked by loxP sites and this the genes are subject to combinatorial deletion when Cre recombinase is induced to enter the nucleus. While these 9 genes are individually all non-essential, their complete deletion would be lethal, making this an essential functional module.

The SGM design used here intentionally preserved the native promoters and terminators for each relocated gene, in recognition of the complex dynamic regulation required for cell cycle oscillations, which is likely partially embedded in the native gene sequence. However, constructing this presented significant challenges. Firstly, the choice of where to start and end each gene was non-obvious, as promoter and terminator lengths vary considerably in yeast and gene relocation can alter expression patterns^34,35^. Our manual curation of each gene’s local DNA sequence was a labour-intensive (and somewhat arbitrary) approach to choosing the DNA fragment to be relocated. Future work would benefit from computational and AI models that predict how the local DNA sequence flanking genes contributes to a gene’s regulation and how local genetic context affects gene expression, especially when genes are moved to new locations^55–57^. Data from synthetic yeast genome SCRaMbLE experiments present a valuable resource for training models to predict expression of genes in new genomic contexts in yeast^58^.

The greater bottleneck for SGM construction was having to delete native genes first before then returning them back to the genome in the growing cluster, which led to having to work with intermediate ‘knock-out’ strains during the process that had slow growth and problems with transformation. Our ‘delete first’ approach was chosen as the alternative ‘build first’ approach would lead to identical copies of the genes being in the genome, which would make it hard to use CRISPR-based methods to delete only the native loci version. However, in retrospect we now recommend that for future SGMs a ‘build first’ strategy would be better. The SGM would need to be assembled from synthetic DNA with versions of genes that are synonymous recoded in key places (e.g. Sc2.0 PCRTags^4^) so that they are not targeted for deletion by the CRISPR systems employed to delete the native gene copies. Further commentary on constructing the cell cycle SGM is given in **Supplementary Note**.

Characterisation of the engineered strains identified phenotypic changes arising from genome reorganisation in our strains. The 6– and 9-gene cluster strains exhibited slower growth rates and delayed cell cycle progression. Expression changes in some of the relocated genes, particularly in *SIC1*, show that gene regulation is not always preserved by relocating a gene with its immediate flanking regulatory elements. Local sequence context influences gene expression and models to predict and design for this are urgently needed^58^. Most critically, we observed aneuploidies in the engineered strains, indicative of genome instability during cluster construction. These may represent an adaptive response to engineering-induced stress^41,59^, a consequence of altered cell cycle gene regulation^60^ or just a side-effect of the strain construction process, which required steps where intermediate strains were created with knock-outs (e.g. Δ*clb2*) known to delay M phase and thus promote aneuploidy generation^42,43^. Regardless of the mechanism, these observations emphasise the technical complexity of simultaneously manipulating multiple cell cycle genes and the importance of regular ploidy monitoring during synthetic genomics projects.

Beyond SGM construction, we also established a novel combinatorial approach for systematic genome minimisation by adapting the SCRaMbLE method to facilitate *in situ* gene deletions. By having estradiol-induced Cre-EBD recombinase target loxP sites placed between cluster genes, we generated hundreds of strains with diverse gene deletion profiles in a single experiment. This approach theoretically allowed us to explore all possible deletion outcomes for the 9-gene cluster simultaneously, rather than laboriously designing and constructing individual strains. To analyse these strains at scale, our POLAR-seq method revealed the frequency of gene loss and retention in minimal strains and determined the most dominant genotypes. POLAR-seq detected approximately 80% of all theoretically possible deletion combinations occurring in our cell pools and proved a cost-effective way to allow us to uncover rare recombination events that would be difficult to identify through individual strain isolation and analysis.

Sequencing revealed that *CLB5* and *SIC1* were the most frequently retained genes aligning with a prior model that predicts these genes are essential components of a minimal cell cycle^23,24,27^. However, we cannot directly equate retention frequency with essentiality without further experiments, such as repositioning these genes within the SGM, at sites where they may be more or less likely to be deleted. We also acknowledge that the altered expression of *SIC1* in our SGM might influence its deletion frequency, potentially confounding our interpretation of its essentiality from the sequencing analysis.

Our data further suggest that just one CLN gene (from the pair of *CLN1,2*), one FKH gene, and *SIC1* are sufficient for sustaining cell growth (assuming CLB genes are present). However, our experimental design had limitations with *CLB1* and *CLB6* not relocated (see **Supplementary Note**). Without their inclusion in the SGM we cannot comprehensively determine the minimal set of CLB genes required for viability. However, despite this limitation, the most abundant genotypes identified in our sequencing closely matched prior models that predict a minimal set consisting of *CLN2*, *CLB5*, *CLB3*, *CLB2*, *SIC1*, and *FKH2* genes^23,24,26,27^. This alignment between our experimental results and theoretical predictions, even with an incomplete set of CLB cyclins in our combinatorial deletion library, provides valuable validation of these computational models.

In summary, we established a novel approach to studying genome minimisation and genetic redundancy by engineering synthetic genome modules for a key set of functionally related genes in *S. cerevisiae*. Through CRISPR-based genome engineering, synthetic gene cluster assembly and subsequent Cre-mediated recombination, we have advanced methods for constructing modular yeast genomes and facilitate better study and control of specific cellular functions like the cell cycle. The SGM combinatorial deletion approach and the use of POLAR-seq to analyse the deletion outcomes has broader applications beyond our cell cycle control case study and represents a powerful methodology for sampling all the gene reduction possibilities for synthetic genome projects.

## Materials and Methods

### Strains and Culture Conditions

#### Yeast strains and media

All yeast strains were derived from *Saccharomyces cerevisiae* BY4741 (*MATa his3Δ1 leu2Δ0 met15Δ0 ura3Δ0*). Strains were cultured in YPD medium (1% (w/v) Bacto Yeast Extract, 2% (w/v) Bacto Peptone, 2% glucose) at 30°C with shaking at 250 rpm unless otherwise stated. Selection was performed on synthetic complete (SC) dropout agar plates containing 2% (w/v) glucose, 0.67% (w/v) Yeast Nitrogen Base without amino acids, 0.14% (w/v) Yeast Synthetic Drop-out Medium Supplements without histidine, leucine, tryptophan, and uracil, supplemented with 20 mg/L tryptophan and 20 g/L bacteriological agar. Depending on selection requirements, media were supplemented with 20 mg/L uracil, 100 mg/L leucine, and 20 mg/L histidine. Alternative carbon sources included YPEtOH (2% ethanol), YPGal (2% galactose), or combinations as specified. For kanMX4 selection, media were supplemented with G-418 disulfate solution to 250 mg ml⁻¹.

#### *E. coli* strains and media

NEB Turbo Competent *E. coli* (F’ *proA^+^B^+^ lacIq ΔlacZM15 / fhuA2 Δ(lac-proAB) glnV galK16 galE15 R(zgb-210::Tn10)Tet^S^ endA1 thi-1 Δ(hsdS-mcrB)5*) were used for cloning and plasmid propagation. Selection and growth were performed in LB medium at 37°C with aeration, supplemented with appropriate antibiotics (ampicillin 100 μg/ml, chloramphenicol 34 μg/ml, or kanamycin 50 μg/ml).

### Molecular Biology

#### DNA manipulation and assembly

DNA parts were constructed using MoClo Yeast Toolkit (YTK) methods^33,61^. PCR amplifications were performed using Phusion-HF or Q5 high-fidelity DNA polymerase (NEB) according to manufacturer’s protocols, using 1 μg genomic DNA or 10 ng plasmid DNA as template in 50 μl reactions. Golden Gate assemblies used equimolar concentrations of 50 fmol/μl DNA fragments with appropriate restriction enzymes (BsaI, BsmBI, or BbsI-HF) in the following thermocycler program: (42°C for 2 min, 16°C for 5 min) × 25 cycles, 60°C for 10 min, 80°C for 10 min. Gibson assemblies were performed at 50°C for 1 hour using in-house master mix with equimolar DNA concentrations of 10 fmol/μl. Small fragment assemblies were created by annealing phosphorylated oligonucleotides with T4 polynucleotide kinase treatment. Primers were obtained from IDT and resuspended to 100 μM stocks. PCR products were purified using DNA Clean & Concentrator Kit (Zymo Research) or gel extractions using Zymoclean™ Gel DNA Recovery Kit. Plasmids were propagated in *E. coli* and purified using Qiagen Miniprep protocols with homemade buffers.

#### Yeast transformations

Chemically competent yeast cells were prepared using the lithium acetate protocol^62^. Overnight YPD cultures were diluted 1:100 and grown to OD_600_ 0.8-1.0, pelleted, washed with 0.1 M LiOAc, and resuspended to 100 μl/transformation. Cell suspensions were mixed with 64 μl DNA/salmon sperm DNA mixture (10 μl boiled salmon sperm DNA + DNA + ddH₂O) and 294 μl PEG/LiOAc mixture (260 μl 50% (w/v) PEG-3350 + 36 μl 1 M LiOAc), heat-shocked at 42°C for 40 min, pelleted, resuspended in 200 μl 5 mM CaCl₂, and plated on selective media.

#### Plasmid curing

For non-URA markers, strains were passaged on non-selective media with serial 1:100 back dilutions for two overnights and confirmed by spotting on selective and non-selective plates. For URA⁺ plasmids, 5-FOA counter selection was used.

### CRISPR-Cas9 Genome Editing

#### Gene Knockout Approaches

Deletion of genes and their flanking regulatory sequences from their native loci in the yeast genome was done via one of two CRISPR-based gene knockout systems. **System One** used two plasmids: a gRNA expression plasmid and a CRISPR/Cas9 expression plasmid (pWS2081 or pWS2083 with URA3 or HIS3 markers). gRNAs were designed to cut between homology arms. For System One, gRNA plasmids were constructed by T4 PNK phosphorylation and annealing of 26 bp oligonucleotides, followed by BsmBI Golden Gate assembly into SpCas9 sgRNA Dropout vector (pWS2069). For in vivo gap repair, 50 ng CRISPR/Cas plasmid and 500 ng each sgRNA plasmid were mixed with 0.5 μl BpiI, 1 μl 10X Buffer G, and water to 10 μl, then incubated at 37°C for 8 hours followed by 80°C inactivation. **System Two** used a single plasmid encoding sgRNA arrays and Cas9. Individual gRNA-tRNA fragments were amplified by Q5 PCR using pWS3178 template (∼2 ng μl⁻¹) and assembled into pWS3910 (URA⁺) by BsaI Golden Gate assembly.

#### Donor DNA design and transformation

Donor DNA contained CRISPR targeting sequences flanked by 500 bp homology arms unless otherwise specified. Donor DNA was PCR-amplified from BY4741 genomic DNA and purified. For transformations, 500 ng each donor DNA fragment was added to CRISPR/gRNA mixtures (10 μl for System One; 250 ng plasmid for System Two) to 64 μl total volume for yeast transformation.

#### Validation of edits

Colonies were screened by colony PCR using Phire Plant Direct PCR Master Mix kit with 1 μl genomic DNA template or colony resuspension. Confirmed edits were verified by Sanger sequencing (Source Bioscience) of PCR-amplified regions spanning ∼100 bp upstream and downstream of edited sites. CRISPR plasmids were cured using 5-FOA counter selection.

#### Whole genome sequencing

Sequencing was performed using Illumina NextSeq 500 with paired-end protocol (MIGS, Pennsylvania, USA). Raw reads were pre-processed with fastp to remove adaptors, then mapped to reference genomes using BWA-MEM2 (Version 2.2.1+galaxy1^63^). Alignments were visualized in Integrative Genomic Viewer (IGV), and coverage was analysed using BAM Coverage Plotter and BAMStats tools on the Galaxy platform.

### Phenotypic Assays

#### Growth assays

For plate reader assays, overnight YPD cultures were back-diluted to OD_600_ 0.2 and grown to mid-exponential phase. Cultures were normalized to OD_600_ 1.0, washed twice with ddH₂O, and diluted 1:100 to starting OD_600_ 0.01 in 96-well plates (100 μl per well, triplicate measurements). Plates were sealed with Breathe-Easy film and incubated in a Synergy HT plate reader at 30°C with shaking, measuring OD_600_ every 15 minutes for 48 hours. For spot assays, cells at OD_600_ 1.0 were serially diluted 1:10 to 10^-5^ in 96-well plates. 5 μl spots were plated on appropriate media and grown for 72 hours static at specified temperatures.

#### Microscopy

Overnight cultures were visualized on a Nikon Eclipse Ti inverted microscope at 20× and 40× magnification. Bright field images were processed using Fiji (ImageJ) to measure cross-sectional areas of 100-300 unbudded single cells using threshold and particle analysis functions.

#### Statistical Analysis

Data are presented as mean ± standard deviation unless otherwise stated. Statistical significance was determined by unpaired t-test using GraphPad Prism software.

### Cell Cycle Analysis

#### Cell synchronization

Mid-exponential cultures (OD_600_ 0.6-0.8) in 100 ml YPD were synchronized by adding 8 μl α-factor (2 mg/ml stock) and incubating for 90 min at 30°C with shaking. Alternatively, hydroxyurea or nocodazole were used for early S phase or G₂/M synchronization. Synchronization efficiency was monitored by budding cell counts. Cultures were washed twice with pre-warmed YPD, resuspended in fresh media, and samples collected every 15 minutes for 90 minutes.

#### Flow cytometry

500 μl culture samples were fixed in 1 ml 90% ethanol at –20°C for ≥18 hours. Fixed cells were pelleted (5000×g, 20 min, 4°C), washed twice with 800 μl 50 mM sodium citrate pH 7.2, and suspended in 500 μl sodium citrate containing 20 μg/ml RNase A and 2.5 μM SYTOX Green. After 1 hour at 37°C, 10 μl proteinase K (20 mg/ml) was added and samples incubated at 55°C for ≥1 hour. Before analysis, 200 μl stained cells were diluted in 2 ml PBS and sonicated (30 sec ON/OFF pulses, 50% amplitude) on ice. DNA content of 100,000 cells was analysed using Attune NxT Flow Cytometer, applying the Watson Pragmatic model in FlowJo software (RRID:SCR_008520) to use fluorescence data to describe cell cycle progression.

#### RT-qPCR

15 ml culture samples were pelleted and snap-frozen. RNA was extracted using NucleoSpin RNA Mini Kit with glass bead disruption supplemented with 1% β-mercaptoethanol. RT-qPCR was performed using Luna Universal One-Step RT-qPCR Kit with 1 μl RNA (100 ng/μL) in 20 μl reactions. Primers were designed using Primer3Plus^64^ and validated for efficiency (R² = 100% ± 15%). Data were normalized against ACT1 (or TAF10 for asynchronous cultures) using the ΔΔCt method^65^. Statistical significance was assessed by unpaired t-test in GraphPad Prism.

### Cre/loxP-Mediated Deletions

#### Induction

Strains transformed with the SCOUT SCRaMbLE reporter plasmid pXL005 (HIS⁺)^29^ were grown to mid-exponential phase and induced with 1 μM β-estradiol (or DMSO control) for 4 hours. Cultures were washed twice, resuspended in 100 μl ddH₂O, serially diluted 10-fold, and 1 μl plated on selective media. Cell viability was assessed by colony counting across plate quadrants after 2-3 days growth at 30°C.

#### FACS sorting

GFP⁺ cells were sorted post-Cre induction using a BD FACSAria III Cell Sorter (70 μm nozzle) after washing and resuspending in PBS. Sorted cells were collected, inoculated into SC LEU⁻HIS⁻ media, or stored in 50% glycerol at –70°C.

#### Multiplex PCR

Multiplex PCR used a single forward primer (AM233, 100 μM, 0.2 μl) targeting loxP sites paired with multiple reverse primers at varying distances. Set 1 tested *CLB5*, *CLB3*, *FKH2*, *CLB2*, *CLN2*, *SIC1* with reverse primers AM234, 236, 237, 238, 390 (0.67 μl each, 10 μM) and AM235 (1 μl, 10 μM). Set 2 tested *LEU2*, *FKH1*, *CLB4*, *CLN1* using reverse primers AM429, 430, 431 (0.67 μl each, 10 μM) and AM432 (1 μl, 10 μM). Reactions contained 1 μl template DNA, 4 μl 5x Herculase Buffer, 0.4 μl dNTPs, 0.6 μl DMSO, 0.2 μl Herculase II polymerase, and water to 20 μl.

### POLAR-seq

The development history and detailed protocol and analysis method for POLAR-seq is available in a separate methods paper^41^. For **High molecular weight DNA isolation,** gDNA was prepared from 100 ml saturated YPD cultures following the Oxford Nanopore protocol with modifications: centrifugation at 4°C, 4000×g for 10 min; 150 μl lyticase (8000 units/ml) for spheroplasting at 30°C; overnight precipitation in TE buffer at 30°C. DNA was purified with AMPure XP beads and quality-assessed by Nanodrop, Qubit, and 0.6% agarose gel. For **long-range PCR and sequencing**, the synthetic gene cluster region was amplified using LA Taq Hot Start (TaKaRa) or repliQa HiFi ToughMix with 10-20 ng gDNA template and 0.1 μM primers in 25 μl reactions. Products were verified on 0.6% agarose gels, purified with AMPure beads, and quantified by Qubit. Libraries were prepared with NEBNext Companion Module and ligation kit SQK-LSK109 or SQK-LSK114, then sequenced using MinION or Flongle flowcells using the MinION Mk1B. For **data analysi**s, reads encompassing the entire module were selected using Porechop for primer identification, excluding reads shorter than the smallest detected amplicon. Genotypes were determined by annotating reads with Liftoff, followed by custom Python script analysis.

## Acknowledgements

We thank Xinyu Lu for help with developing SCRaMbLE selection systems used in the research. This research was supported by a Darwin Trust of Edinburgh PhD scholarship to A.M., a Wellcome Trust Discretionary Award (221267/Z/20/Z) providing funding for K.C. and T.E., a UKRI EPSRC award (EP/M002306/1) providing funding for W.M.S. and T.E., and the Systems Biology Grant of the University of Surrey to M.B.

## Author Contributions

Conceptualised the study: M.B. and W.M.S. Designed and initiated the study: A.M., M.B. W.M.S. and T.E. Performed experiments: A.M. and L.C.A. with help from W.M.S. and K.C. Analysed the data: A.M., L.C.A., K.C., A.G., W.M.S., M.B. and T.E. Supervised the project: M.B. and T.E. Wrote the manuscript: A.M., A.G. and T.E. with input from all authors.

## Competing Interest Statement

K.C. is currently employed by Oxford Nanopore Technologies but was exclusively employed by Imperial College London at the time of data collection for this study. All other authors declare no conflicts of interest.

## Supplementary Materials File

**Figure S1.**
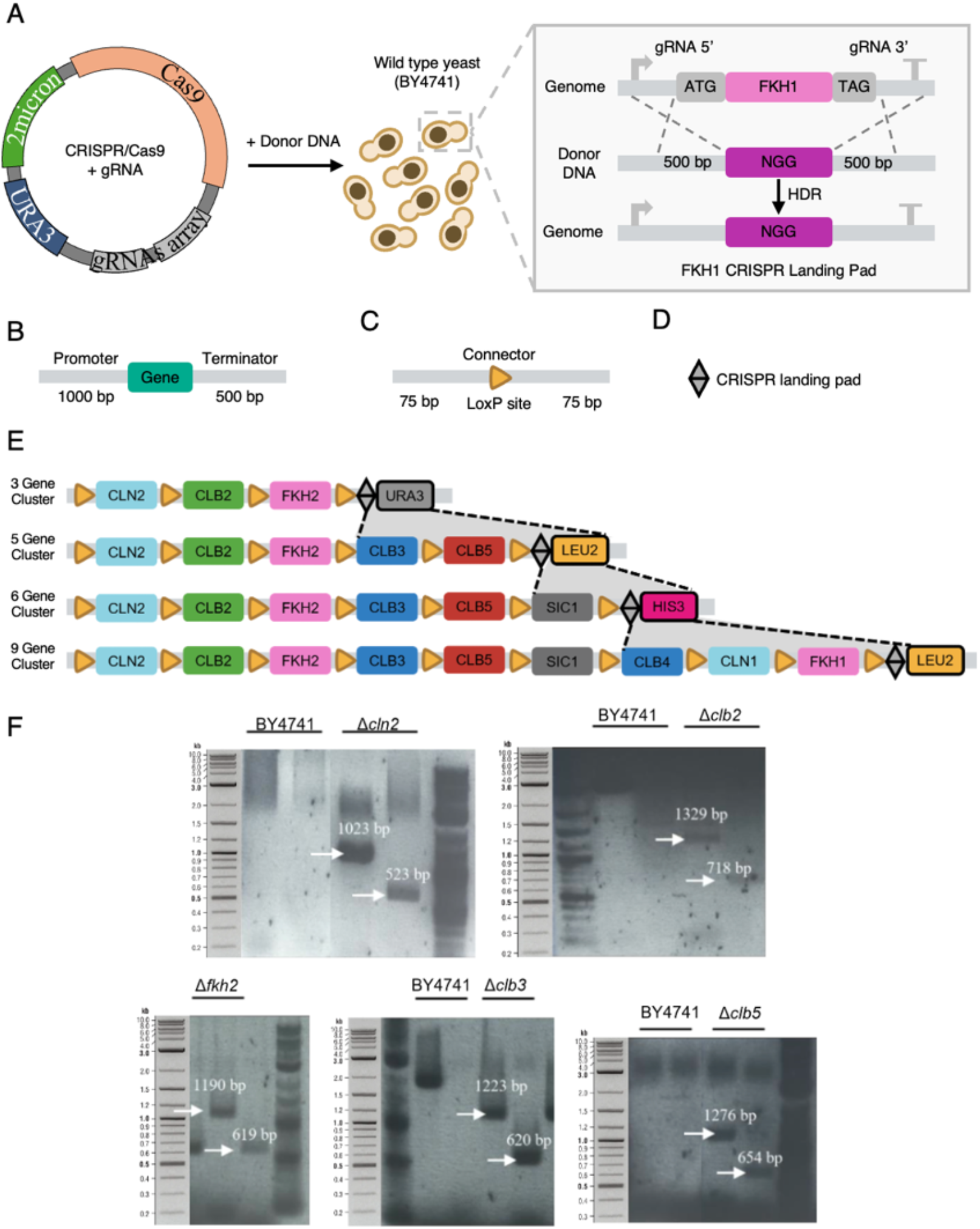
CRISPR/Cas9 Landing Pad Methodology: **(A)** CRISPR/Cas9-mediated approach for ORF deletion and landing pad integration. Wild-type BY4741 yeast cells are transformed with CRISPR/Cas9 plasmid encoding guide RNAs targeting the 5’ and 3’ regions of the target gene (example shown: *FKH1*). Two guide RNAs create double-strand breaks flanking the open reading frame. Homology-directed repair using provided donor DNA replaces the deleted gene with a 24 bp synthetic landing pad sequence containing a unique 20 bp spacer and functional NGG PAM site, flanked by 500 bp homology arms. The resulting edited genome contains the landing pad sequence that can be targeted for future modifications or gene restoration. HDR, homology-directed repair. **(B)** Schematic of gene fragment design showing the standard approach for retaining regulatory elements. Each gene fragment includes ∼1000 bp promoter region (grey), the complete gene sequence (teal), and ∼500 bp terminator region (grey) to preserve native transcriptional regulation when relocated to the synthetic cluster. **(C)** Connector sequence design containing loxP sites (orange triangles), flanked by unique 75 bp sequences for homology-directed assembly. **(D)** CRISPR landing pad (black diamond) positioned at the 3’ end of each cluster iteration to enable targeted cluster expansion in subsequent gene editing rounds. **(E)** Stepwise cluster expansion showing the iterative assembly from 3-gene to 9-gene clusters. Each cluster maintains consistent gene orientation (5’ to 3’) with loxP-flanked connectors between genes. Auxotrophic selection markers (*URA3*, *LEU2*, *HIS3*) are swapped during each expansion round: 3-gene cluster with *URA3* (grey), 5-gene cluster with *LEU2* (yellow), 6-gene cluster with *HIS3* (magenta), and 9-gene cluster with *LEU2* (yellow). The CRISPR landing pad (black diamond) enables targeting for each successive expansion round. **(F)** Representative agarose gel images showing individual gene knockout verification (Δ*cln2*, Δ*clb2*, Δ*fkh2*, Δ*clb3*, Δ*clb5*). Yeast transformants that grew on selective media were checked by colony PCR using two unique primer sets for each gene. The first set of primers targeted the integration region outside the homology arms (HA), while the second set amplified the region from outside the 5’ homology arm into the landing pad (LP). 1% agarose gel electrophoresis was performed with NEB Quick-Load® 1 kb Plus DNA Ladder. Black arrows indicate PCR products corresponding to successful gene deletions, with specific band sizes shown (e.g., 1023 bp and 523 bp for Δ*cln2*; 1329 bp and 718 bp for Δ*clb2*). BY4741 serves as the wild-type control.

**Figure S2.**
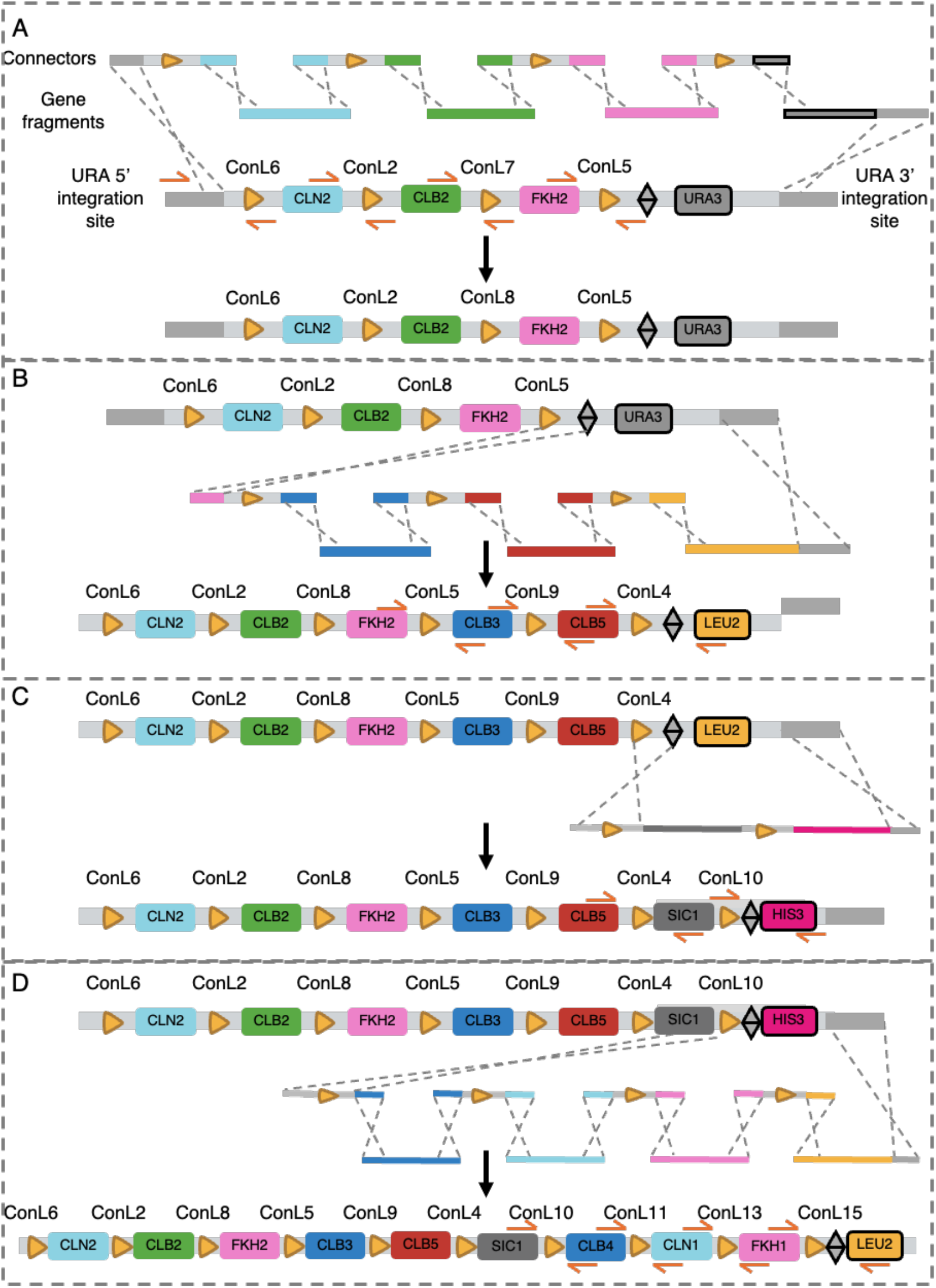
Detailed assembly of the cluster versions from DNA parts: **(A)** The three gene cluster was first assembled into the URA locus by homology-based assembly of 3 PCR-amplified gene fragments, the URA3 selectable marker gene and four synthetic connectors, each containing LoxP sites (triangle). **(B)** The five gene cluster was constructed by expansion of the three gene cluster and URA3 to LEU2 marker exchange. **(C)** The six gene cluster added in the SIC1 gene fragment and exchanged the marker to HIS3. **(D)** The nine gene cluster added 3 more gene fragments assembled with synthetic connectors and returned to the LEU2 selectable marker.

**Figure S3:**
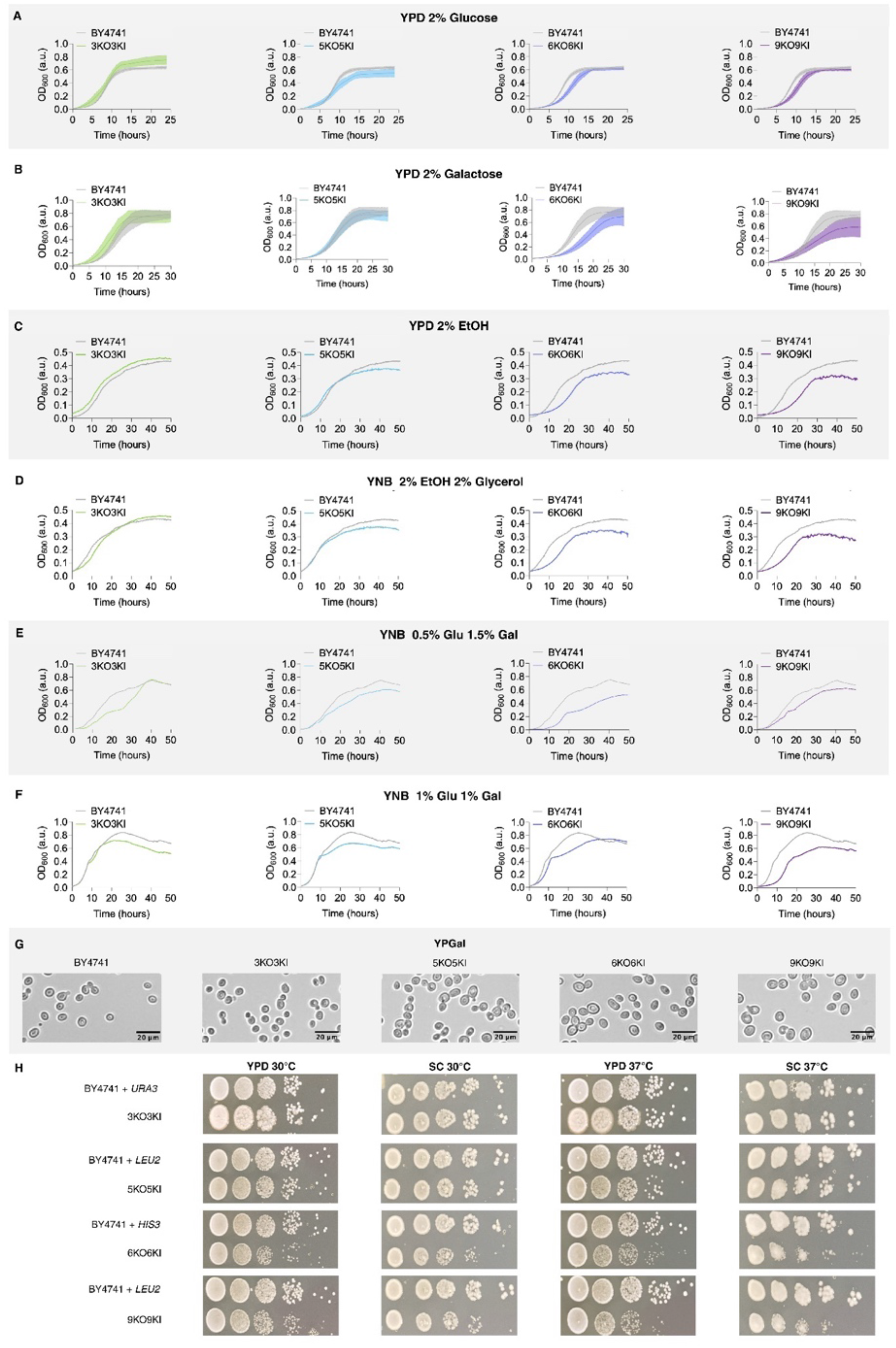
Phenotypic characterisation of synthetic cell cycle cluster strains across diverse growth conditions. Growth and viability of the 3-, 5-, 6-, and 9-gene synthetic cell cycle cluster strains. **(A–F)** Growth curves measured by plate reader assay (OD_600_) in various media at 30°C. **(A)** YPD 2% glucose and **(B)** YP 2% galactose (mean ± SEM of 5 and 4 biological replicates, respectively; error bars shown as dotted lines and semi-transparent fill areas). **(C–F)** Non-standard carbon sources tested with single biological replicates: **(C)** YNB 2% ethanol, **(D)** YNB 2% ethanol + 2% glycerol, **(E)** YNB 0.5% glucose + 1.5% galactose, and **(F)** YNB 1% glucose + 1% galactose. For panels **A–C**, seed cultures were grown in the same media used for growth measurement. For **D–F**, cultures were grown in YPD 2% glucose, washed twice in YNB without carbon source, then inoculated into measurement media. Controls: yAM075 (BY4741 with *LEU2* reintegrated at the *URA3* locus) for YPD 2% glucose condition; yAM017 (BY4741 with *HIS3* reintegrated at *URA3*) for all other conditions. **(G)** Representative bright-field microscopy images of WT and synthetic cluster strains grown in YP 2% galactose (20× mag.; scale bar = 20 μm). **(H)** Spot assay viability of WT and synthetic cluster strains on 2% glucose (YPD and SC media) grown at 30°C or 37°C. Serial 10-fold dilutions from normalised cultures (OD_600_ = 1.0) were spotted from left to right and incubated for 3 days (n=3 for 9-gene cluster and yAM075).

**Figure S4:**
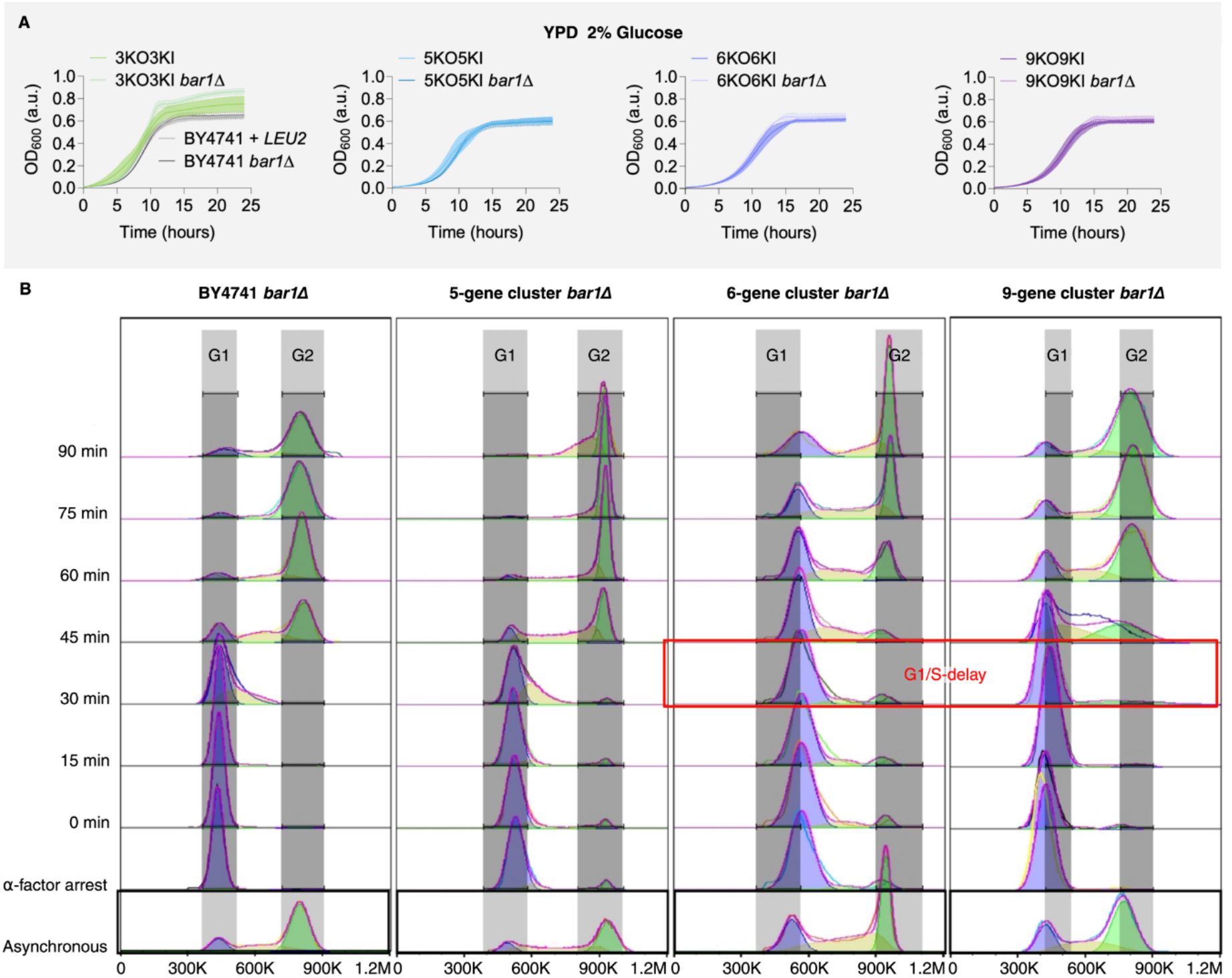
Alpha-factor synchronisation and cell cycle progression analysis of *bar1Δ* synthetic cluster strains. **(A)** Growth comparison of *BAR1* and *bar1Δ* strains in YPD 2% glucose at 30°C. Mean OD_600_ ± SEM from 5 (*BAR1*) and 2 (*bar1Δ*) biological replicates; error bars shown as dotted lines with semi-transparent fill areas. **(B)** Cell cycle progression of α-factor synchronised *bar1Δ* strains. Cells arrested at G_1_/S (START) were released and sampled at 15-minute intervals (0–90 min) from BY4741 *bar1Δ* control and *bar1Δ* versions of 5-, 6-, and 9-gene cluster strains (n=2). Flow cytometric histograms with SYTOX Green staining shown as half-offset overlays. Asynchronous controls (black frames) were used to define G_1_ and G_2_ gates (grey boxes). Watson Pragmatic cell cycle modelling of SYTOX Green fluorescence intensity was performed in FlowJo to determine G_1_ (blue), S (yellow), and G_2_/M (green) phase cells. Red box indicates G_1_/S-phase delay in 6– and 9-gene strains (60 and 45 min, respectively).

**Figure S5:**
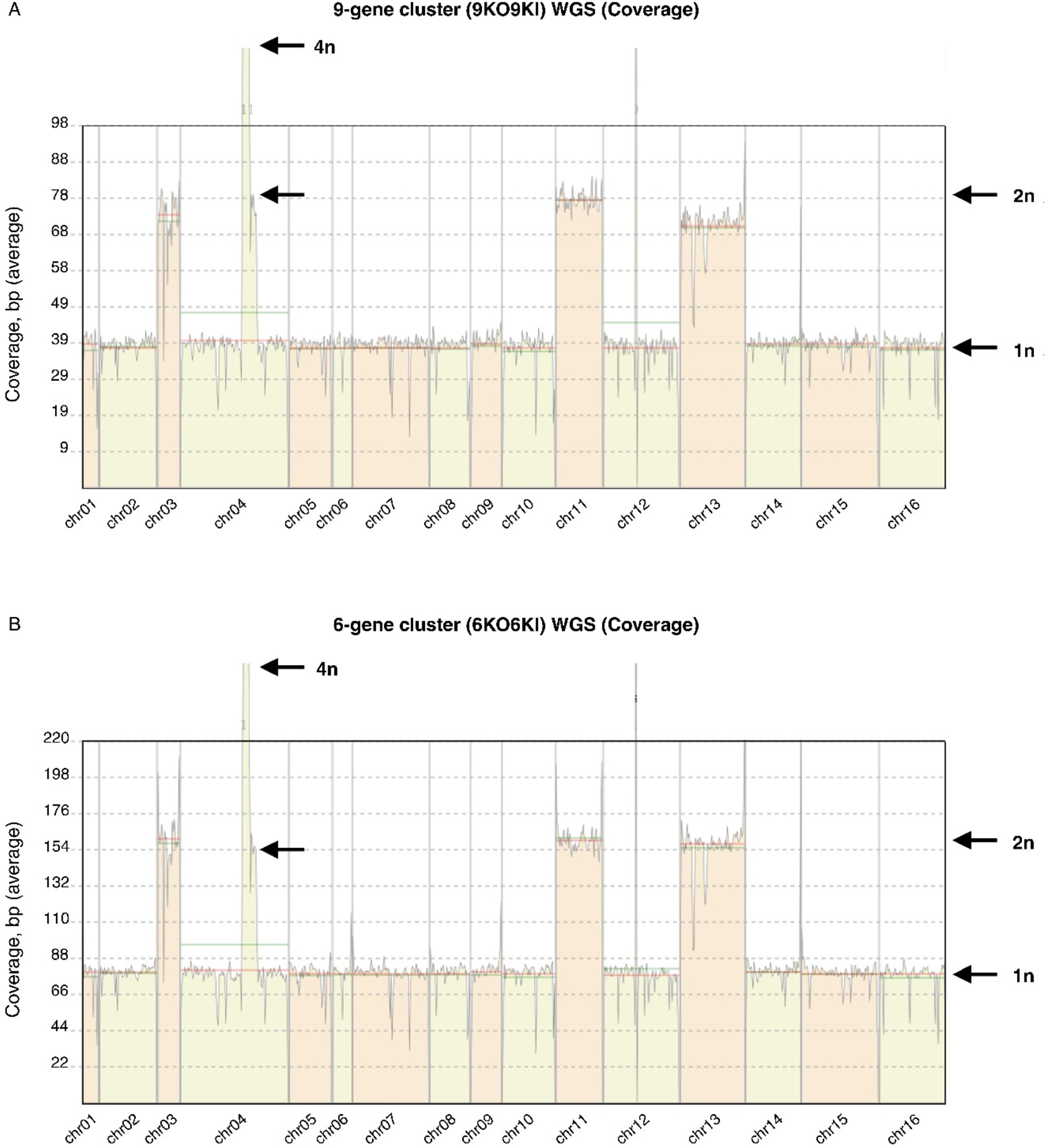
Whole genome sequencing reveals chromosome aneuploidies in 6– and 9-gene cluster strains. Whole genome sequencing (WGS) coverage analysis reveals chromosome aneuploidies in synthetic cluster strains. Read depth across all chromosomes in **(A)** 9-gene cluster and **(B)** 6-gene cluster strains. Data visualised using BAM Coverage Plotter. Reference coverage (1n) and duplicated regions (2n and 4n) are indicated by arrows. Both strains show 2n coverage on chromosomes III, XI, and XIII, plus additional 4n (115 kb, chr IV) and 2n (109 kb, chr IV) duplications in chromosome IV, collectively representing ∼18% of genome duplication based on the length of these chromosomes.

**Figure S6:**
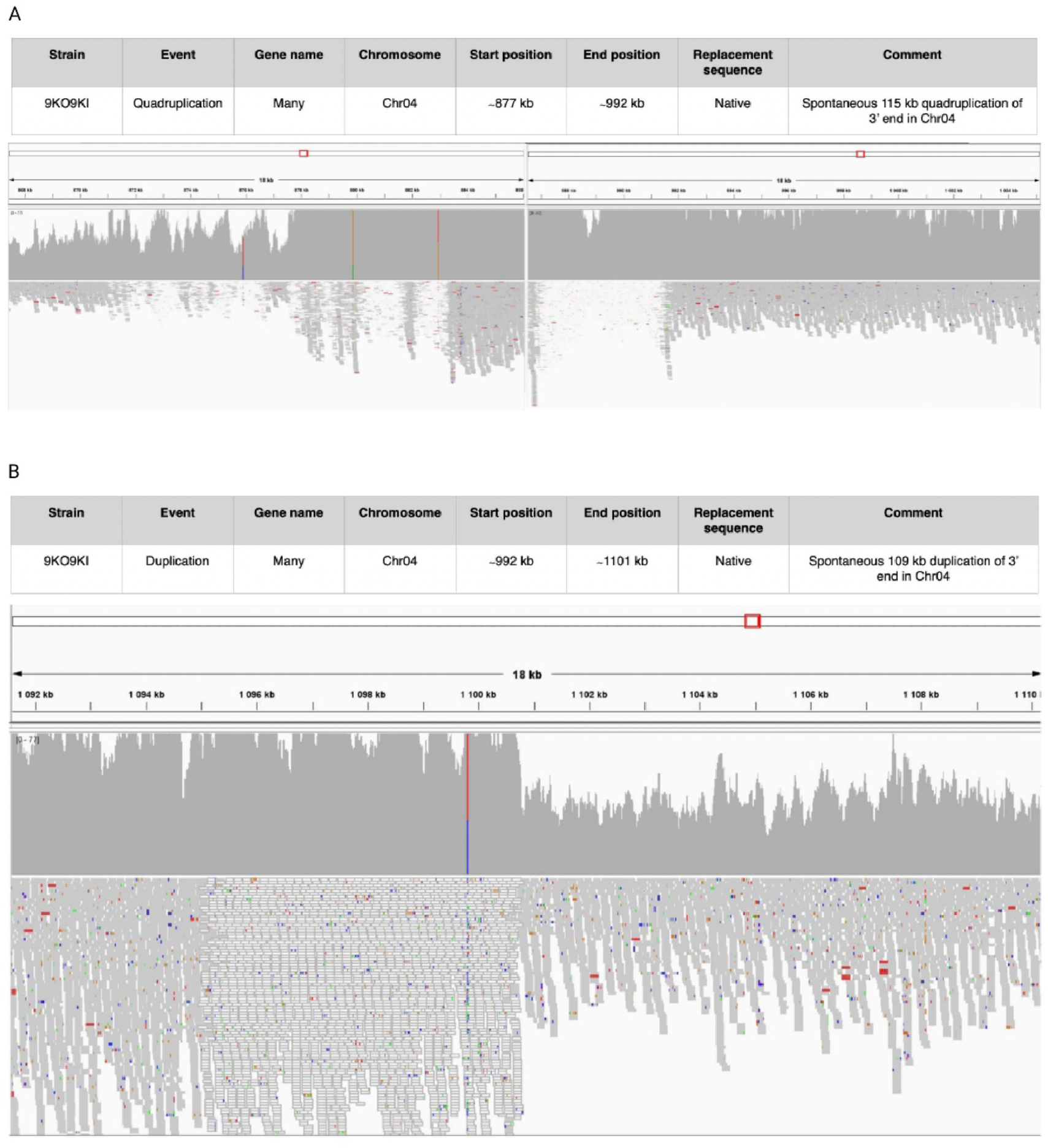
Detailed analysis of chromosome IV rearrangements in 9-gene cluster strain by whole genome sequencing. Integrative Genomics Viewer (IGV) visualisation of chromosome IV rearrangements in 9-gene cluster strain. **(A)** Coverage distribution (grey histogram, top) and read alignment (bottom) showing the transition from single (1n) to quadruple (4n) coverage at position ∼877–992 kb (115 kb quadruplication). **(B)** IGV coverage map (grey histogram, top) and read alignment (bottom, coloured reads) displaying the junction where 4n coverage transitions to double (2n) coverage (∼992–1101 kb, 109 kb duplication). Genomic coordinates of rearrangement boundaries are annotated in the data tables above each panel.

**Figure S7:**
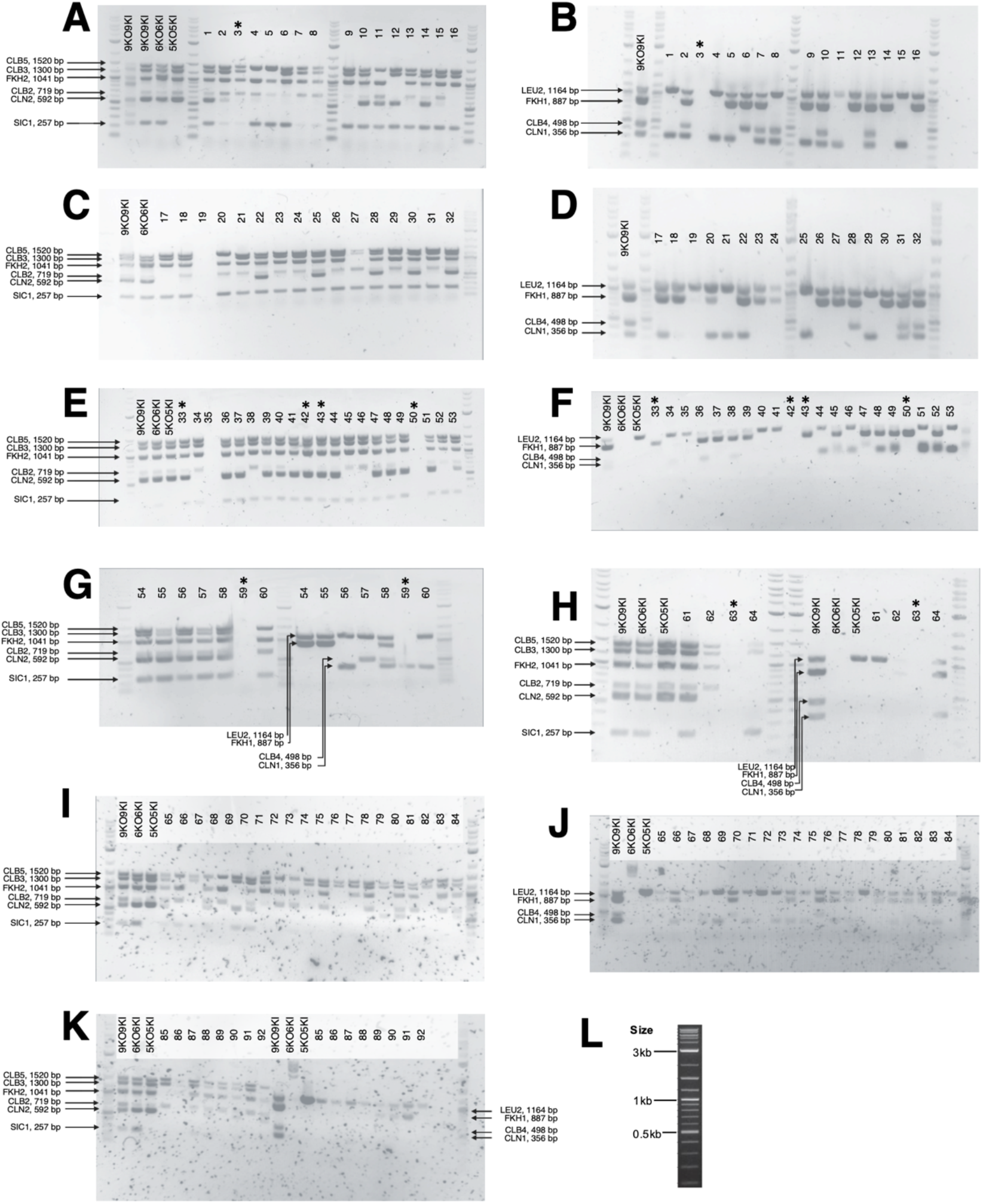
Multiplex PCR-based genotyping of *in vivo* combinatorially minimised synthetic cluster variants. Multiplex PCR screening of 92 *in vivo* deletion colonies reveals gene deletion patterns in synthetic cell cycle cluster variants. Agarose gel electrophoresis (1%, SYBR-Safe staining, visualised with Bio-Rad GelDoc) of multiplex PCR amplicons from individual colonies (numbered across top). Expected band sizes for each target gene are indicated on the left margin. Absence of a band indicates deletion of that gene from the cluster. **Controls:** Parental cluster DNA (9-, 6-, and 5-gene cluster parental strains shown at left of each gel) serves as positive controls to confirm expected band sizes for intact genes. **Set 1 (A, C, E, G left, H left, I, K left):** PCR reactions amplifying *CLB5*, *CLB3*, *FKH2*, *CLB2*, *CLN2*, and *SIC1* using forward primer AM233 and reverse primers AM234–238 and AM390. **Set 2 (B, D, F, G right, H right, J, K right):** PCR reactions amplifying *LEU2*, *FKH1*, *CLB4*, and *CLN1* using forward primer AM233 and reverse primers AM429–432. (**L**) ladder used in all gels. Asterisks (*) denote strains discarded after no *LEU2* amplification was seen in the multiplex PCR.

**Figure S8:**
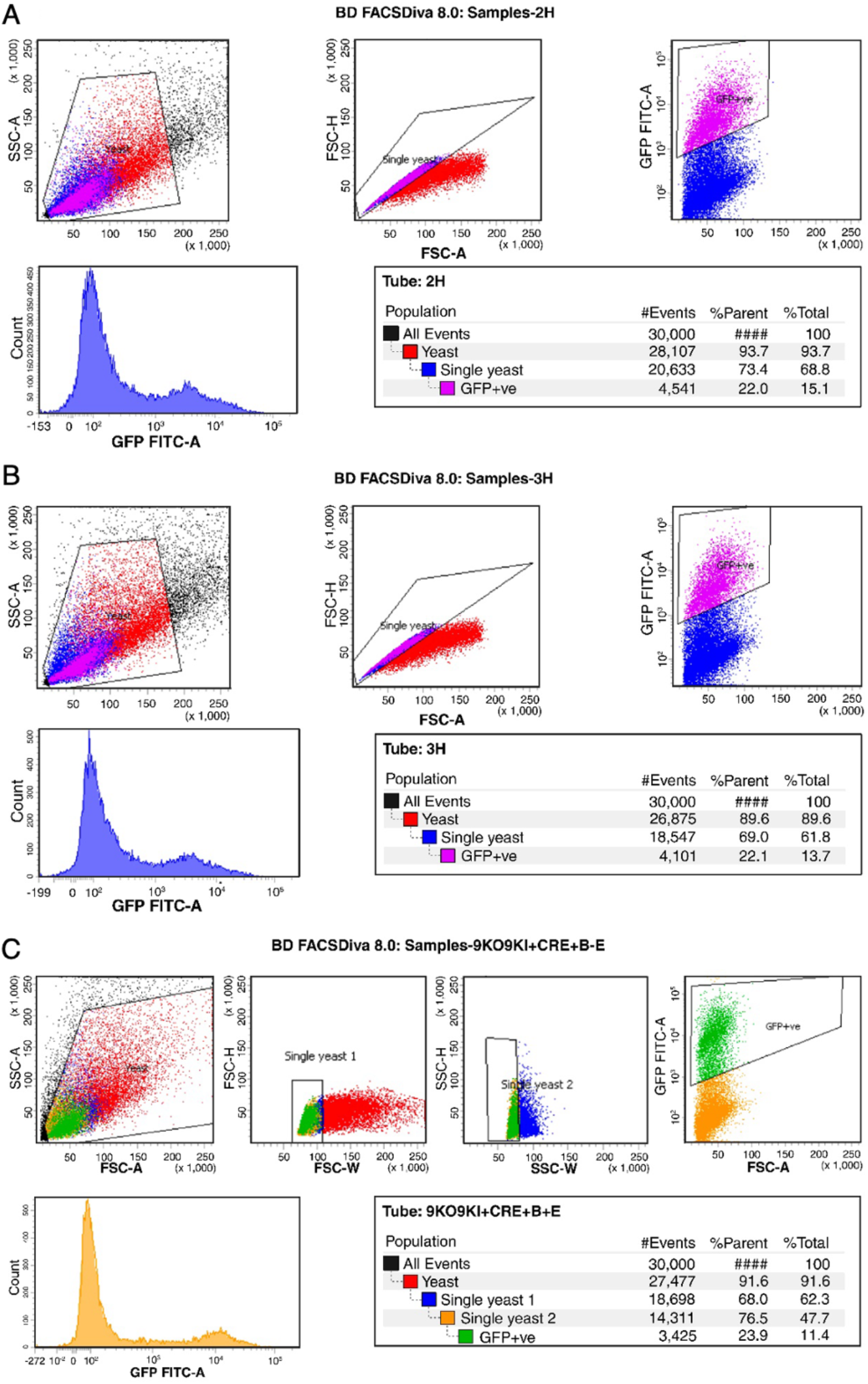
FACS-based isolation of GFP-positive cells from Cre/loxP-recombined synthetic cluster strains. Data acquisition and analysis were performed by Imperial College London Flow Cytometry Facility staff using BD FACSDiva™ Software v8.0 on a BD FACSAria III Cell Sorter. **(A)** Representative 2-hour β-estradiol induction (single biological replicate; Tube: 2H). GFP-positive population: 15.1% of total cells, 22.0% of single yeast. **(B)** First biological replicate of 3-hour β-estradiol induction (Tube: 3H); GFP-positive population: 13.7% of total, 22.1% of single yeast. **(C)** Second biological replicate of 3-hour induction; GFP-positive population: 11.4% of total, 23.9% of single yeast. Sequential gating strategy identified yeast cells (FSC-A vs. SSC-A), single cells to exclude doublets (FSC-A vs. FSC-H), and GFP-positive cells (GFP FITC-A intensity). Panel **(C)** includes two additional refinement gates: FSC-H vs. FSC-W and SSC-H vs. SSC-W to further exclude aggregates. Histogram showing GFP FITC-A distribution indicates low but consistent GFP expression in sorted populations across induction conditions.

**Figure S9:**
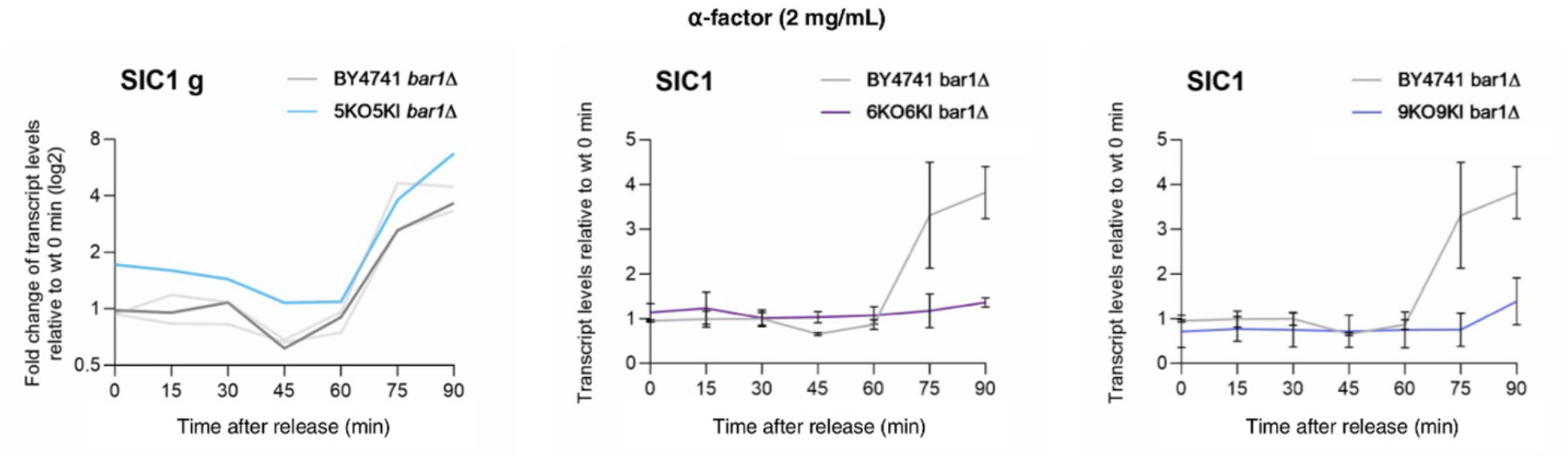
qPCR analysis of *SIC1* gene expression following α-factor synchronisation and release. *SIC1* transcript levels measured by qPCR in α-factor-synchronised *bar1Δ* cells during a single cell cycle following release. Cells were arrested in G_1_ phase by addition of α-factor (2 mg/mL) and incubated for 90 min at 30°C with 250 rpm agitation; synchronisation efficiency was monitored by number of budding cells under light microscopy. After washing twice with pre-warmed YPD medium and centrifugation, cultures were released into warm fresh rich media at 0 min (start of synchronous cell cycle progression). Samples were collected at 15-minute intervals (0–90 min post-release) for RNA extraction and qPCR. *SIC1* transcript levels were quantified using qPCR with the Ct value method, normalised against *ACT1* as the reference gene. *SIC1* transcript fold change is displayed for three synthetic cluster strains — *bar1Δ* 5-gene (aqua blue), 6-gene (indigo), and 9-gene (purple) strains — each compared to the *bar1Δ* wild-type control (dark grey line) in the left, middle, and right panels. Error bars represent standard deviation from independent biological replicates: wild-type *bar1Δ* (n=3), 5-gene cluster (n=1), 6-gene cluster (n=2), and 9-gene cluster (n=3).

**Figure S10:**
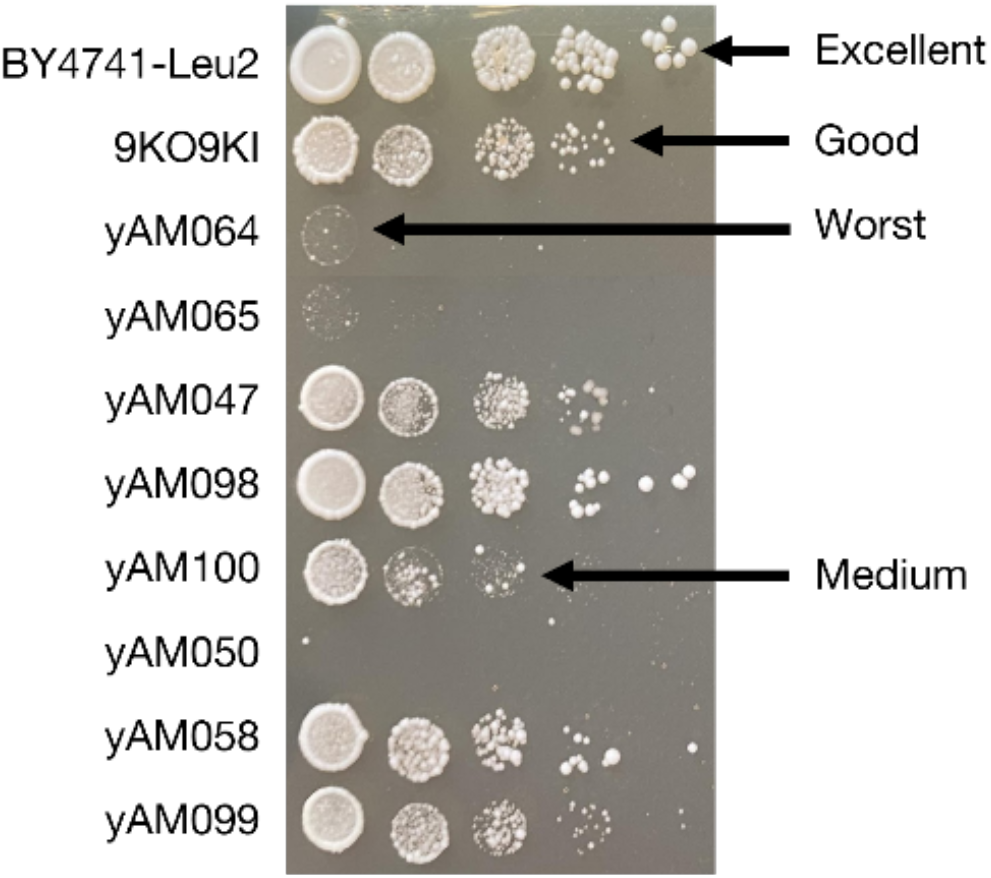
Method for spot assay-based quantification of relative viability. Spot assay in YPD 37◦C: Each column is a different OD_600_ dilution starting at OD_600_ = 1 with 1:10 dilutions steps (n = 2). Arrows indicate how spot assays were numerically quantified to assess relative viability following the guidelines that (i) a score of 1 (Excellent) to strains that had number of colonies and colony size similar to BY4741-Leu2, (ii) a score of 2 (Good) to strains that had number of colonies and size similar to 9KO9KI, (iii) a score of 3 (Medium) to strains that had no colonies in the lowest dilution that the 9KO9KI displayed colonies, (iv) a score of 4 (Bad) to strains that had no colonies in at least two dilution steps higher than the lowest dilution the 9KO9KI strain showed colonies, and (v) a score of 5 (Worst) to strains that showed no colonies or barely any colonies in OD_600_ 1 spot.

**Figure S11:**
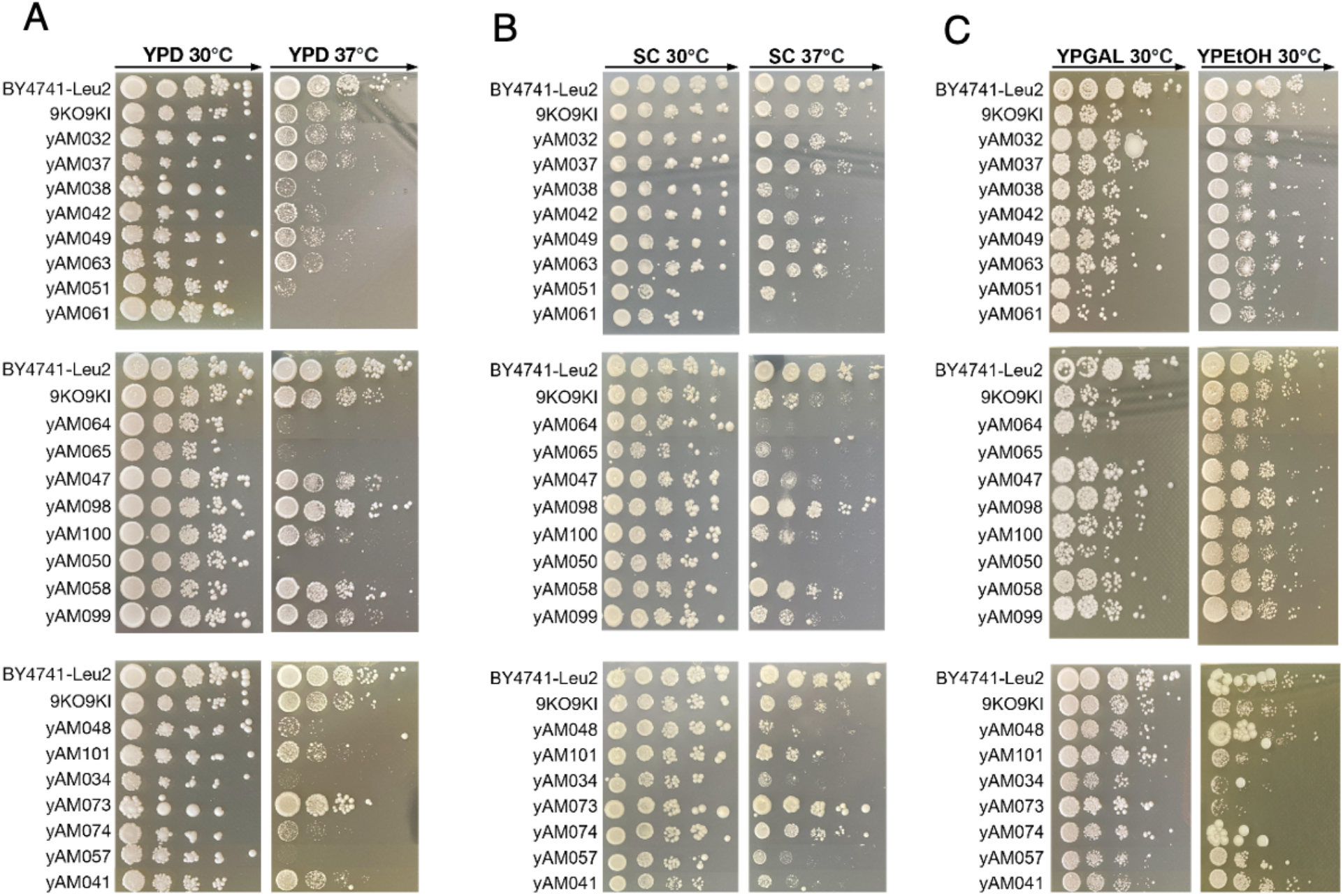
Spot assay of minimal viable cell cycle strains cells on different carbon sources. Spot assays under different conditions. Ten-fold serial dilutions of normalised cultures (OD_600_ = 1.0) were spotted from left to right on media with different carbon sources: 2% glucose (YPD and SC media, panels (A) and (B) and incubated at 30°C or 37°C for 3 days. n=3 for the 9-gene cluster and the control strain yAM075 (BY4741 with *LEU2* reintegrated at the *URA3* locus). (C) YPGal (2% Galactose) and 30◦C YP-EtOH (2% Ethanol) incubated at 30°C or 37°C for 3 days.

**Figure S12:**
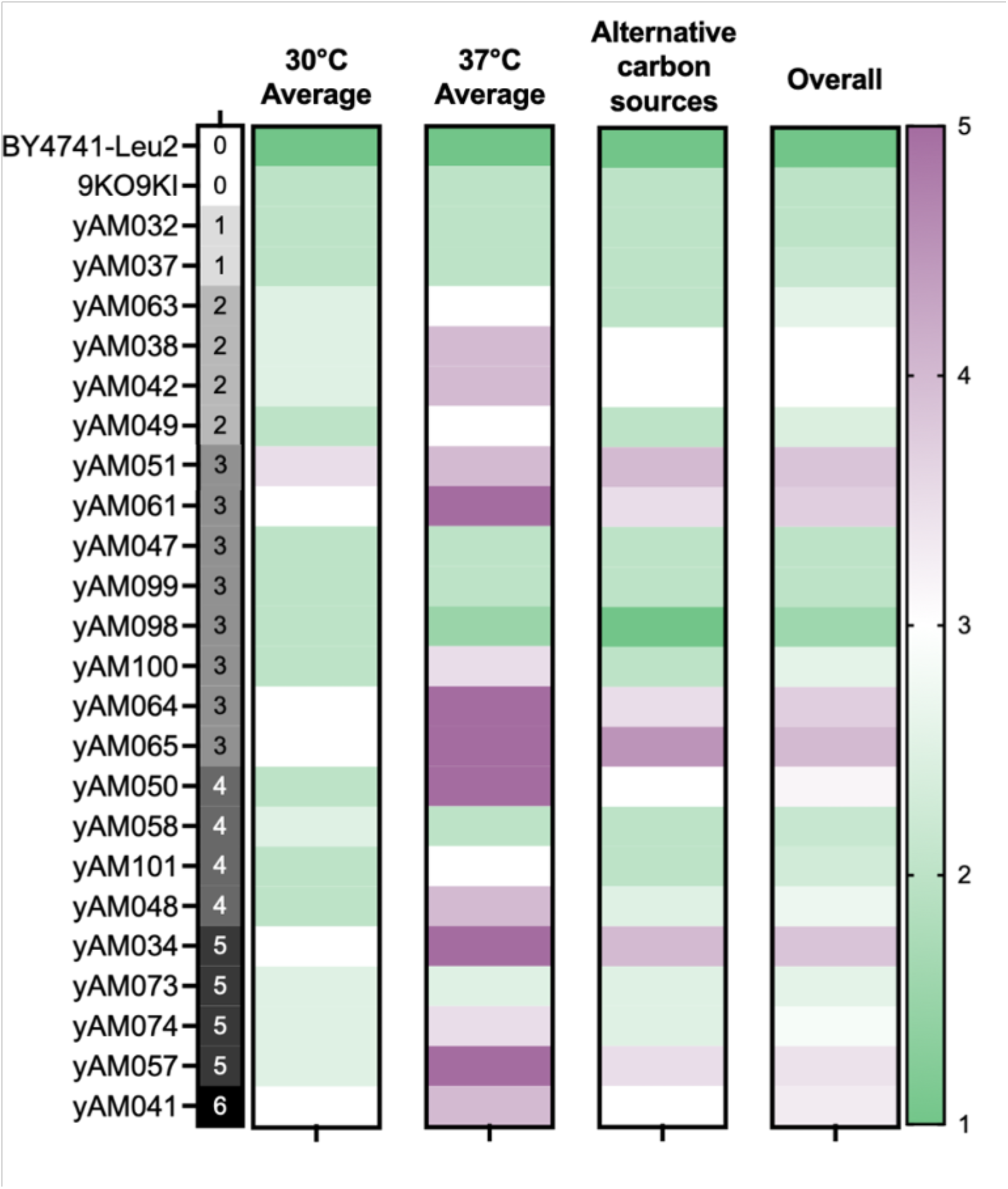
Heatmap to represent average solid-phase growth performance of the strains in different conditions. Spot assay data from Figure S11 was qualitatively scored following the guide in Figure S10. Then for each strain the average score was determined for growth at 30°C (on YPD and SC-Glu media), at 37°C (on YPD and SC-Glu media) and on alternative carbon sources (YP-Gal and YP-EtOH at 30°C). An overall score was also determined by averaging across all conditions for each strain. Values were then shaded with 1 (excellent growth) as green, 3 (medium performance) as white and 5 (worst performance) as purple.

**Table S1:** Complete list of the unique genotypes discovered from MinION sequencing of FACS-sorted post-Cre/loxP cell pool. Full list of genotypes from MinION. The table shows an ordered list of unique cluster strains, the cluster content for each strain, the number of reads corresponding to each genotype in the ‘count’ column and the percent of each genotype in the MinION sequencing run. Parental genotype (combination 29) is highlighted in peach.

|  | Genotype | Count | Percent |
| --- | --- | --- | --- |
| 0 | CLN2_CLB3_CLB5_SIC1_FKH1_LEU2 | 40528 | 16.50% |
| 1 | CLN2_CLB2_FKH2_CLB3_CLB5_SIC1_LEU2 | 38375 | 15.60% |
| 2 | CLB2_FKH2_CLB3_CLB5_SIC1_CLN1_LEU2 | 33650 | 13.70% |
| 3 | CLN2_CLB2_FKH2_CLB3_CLB5_SIC1_FKH1_LEU2 | 13265 | 5.40% |
| 4 | CLN2_CLB2_FKH2_CLB3_CLB5_SIC1_CLB4_FKH1_LEU2 | 11866 | 4.80% |
| 5 | CLB2_FKH2_CLB5_SIC1_CLN1_LEU2 | 11621 | 4.70% |
| 6 | CLB2_FKH2_CLB3_CLB5_SIC1_CLN1_FKH1_LEU2 | 11302 | 4.60% |
| 7 | CLN2_CLB3_CLB5_SIC1_CLB4_FKH1_LEU2 | 8115 | 3.30% |
| 8 | CLB3_CLB5_SIC1_CLN1_FKH1_LEU2 | 5825 | 2.40% |
| 9 | CLB2_FKH2_CLB5_SIC1_CLB4_CLN1_FKH1_LEU2 | 5607 | 2.30% |
| 10 | CLN2_CLB2_FKH2_CLB3_CLB5_SIC1_CLB4_CLN1_LEU2 | 5598 | 2.30% |
| 11 | CLB3_CLB5_SIC1_CLN1_LEU2 | 4540 | 1.80% |
| 12 | CLB2_FKH2_CLB3_CLB5_SIC1_CLB4_CLN1_LEU2 | 3782 | 1.50% |
| 13 | CLN2_CLB3_CLB2_FKH2_CLB3_CLB5_SIC1_LEU2 | 3371 | 1.40% |
| 14 | CLB2_FKH2_CLB3_CLB5_SIC1_LEU2 | 3197 | 1.30% |
| 15 | CLN2_CLB4_CLB3_CLB5_SIC1_CLB2_FKH2_FKH1_LEU2 | 2965 | 1.20% |
| 16 | CLB3_CLB5_SIC1_CLN2_LEU2 | 2663 | 1.10% |
| 17 | CLB3_CLN2_FKH2_CLB3_CLB5_SIC1_LEU2 | 2602 | 1.10% |
| 18 | CLN2_CLB3_CLB5_SIC1_LEU2 | 2078 | 0.85% |
| 19 | CLB5_SIC1_CLN1_FKH1_LEU2 | 1992 | 0.81% |
| 20 | CLN2_CLB2_FKH2_CLB3_CLB5_SIC1_CLN1_FKH1_LEU2 | 1969 | 0.80% |
| 21 | CLB2_CLB5_SIC1_FKH1_LEU2 | 1803 | 0.73% |
| 22 | CLN1_CLB3_CLB5_SIC1_LEU2 | 1669 | 0.68% |
| 23 | CLN2_CLB3_CLB5_SIC1_CLB4_CLN1_FKH1_LEU2 | 1546 | 0.63% |
| 24 | CLN2_CLB2_CLN2_CLB5_SIC1_FKH1_LEU2 | 1531 | 0.62% |
| 25 | CLN2_CLB5_SIC1_FKH1_LEU2 | 1508 | 0.61% |
| 26 | CLN2_CLB2_FKH2_CLB3_CLB5_SIC1_CLN1_LEU2 | 1504 | 0.61% |
| 27 | CLB3_CLB5_SIC1_CLN2_CLB4_FKH1_LEU2 | 1385 | 0.56% |
| 28 | CLB5_CLB4_CLN2_CLB2_LEU2 | 1318 | 0.54% |
| 29 | CLN2_CLB2_FKH2_CLB3_CLB5_SIC1_CLB4_CLN1_FKH1_LEU2 | 1257 | 0.51% |
| 30 | CLB3_CLB5_CLB2_FKH2_FKH1_LEU2 | 1187 | 0.48% |
| 31 | CLB2_FKH2_CLB3_CLB5_SIC1_FKH1_LEU2 | 1131 | 0.46% |
| 32 | CLN2_CLB2_FKH2_CLB3_CLB5_SIC1_CLB4_SIC1_CLB4_FKH1_LEU2 | 982 | 0.40% |
| 33 | CLB2_FKH2_CLB3_CLB5_SIC1_CLB4_FKH1_LEU2 | 867 | 0.35% |
| 34 | CLB3_CLB5_SIC1_CLB4_CLN1_LEU2 | 823 | 0.33% |
| 35 | CLB2_FKH2_CLB5_LEU2 | 801 | 0.33% |
| 36 | CLN2_CLB3_CLB5_FKH1_LEU2 | 661 | 0.27% |
| 37 | CLN2_CLB3_CLB5_SIC1_CLN1_FKH1_LEU2 | 614 | 0.25% |
| 38 | CLB3_CLB5_SIC1_FKH1_LEU2 | 606 | 0.25% |
| 39 | CLB2_FKH2_FKH1_LEU2 | 559 | 0.23% |
| 40 | CLN2_CLB3_CLB5_SIC1_CLN1_LEU2 | 543 | 0.22% |
| 41 | CLN2_CLB2_FKH2_FKH1_LEU2 | 501 | 0.20% |
| 42 | SIC1_CLB3_CLN1_CLB5_FKH2_LEU2 | 446 | 0.18% |
| 43 | FKH2_CLB3_CLB5_SIC1_CLB3_CLB5_SIC1_CLN1_FKH1_LEU2 | 380 | 0.15% |
| 44 | CLN1_CLB3_CLB5_SIC1_FKH1_LEU2 | 353 | 0.14% |
| 45 | CLB2_CLB3_CLB5_SIC1_CLB4_CLN1_LEU2 | 338 | 0.14% |
| 46 | CLB2_FKH2_CLB5_SIC1_LEU2 | 328 | 0.13% |
| 47 | CLB2_FKH2_CLB3_CLB5_LEU2 | 298 | 0.12% |
| 48 | CLN1_FKH1_CLB3_CLB5_SIC1_LEU2 | 292 | 0.12% |
| 49 | CLB2_FKH2_CLB5_CLN1_LEU2 | 254 | 0.10% |
| 50 | CLB3_CLB5_SIC1_CLB4_FKH1_LEU2 | 209 | 0.08% |
| 51 | CLN2_CLB3_CLB5_SIC1_CLB4_CLN1_LEU2 | 196 | 0.08% |
| 52 | CLN2_CLB2_FKH2_CLB3_CLB5_SIC1_CLB3_CLB5_SIC1_CLB4_LEU2 | 185 | 0.08% |
| 53 | FKH2_CLN1_CLB3_CLB5_SIC1_LEU2 | 184 | 0.07% |
| 54 | CLN2_CLB3_CLB5_SIC1_CLB3_CLB5_SIC1_FKH1_LEU2 | 182 | 0.07% |
| 55 | CLB2_FKH2_CLB5_SIC1_FKH1_LEU2 | 177 | 0.07% |
| 56 | CLB2_FKH2_CLB5_SIC1_CLN1_FKH1_LEU2 | 176 | 0.07% |
| 57 | CLB2_FKH2_CLB5_SIC1_CLB4_FKH1_LEU2 | 162 | 0.07% |
| 58 | FKH2_CLB3_CLB5_SIC1_LEU2 | 159 | 0.06% |
| 59 | CLN2_CLB2_FKH2_CLB5_SIC1_CLN1_LEU2 | 155 | 0.06% |
| 60 | CLB2_FKH2_CLB5_SIC1_CLB4_CLN1_LEU2 | 152 | 0.06% |
| 61 | CLN1_CLB3_CLB5_SIC1_CLN1_CLB3_CLB5_SIC1_FKH1_LEU2 | 144 | 0.06% |
| 62 | CLB2_FKH2_CLB3_CLB5_SIC1_CLB4_CLN1_FKH1_LEU2 | 143 | 0.06% |
| 63 | CLN2_CLB4_CLB3_CLB5_SIC1_LEU2 | 133 | 0.05% |
| 64 | CLN2_CLB4_CLB3_CLB5_SIC1_FKH1_LEU2 | 88 | 0.04% |
| 65 | CLN2_CLB2_FKH2_CLB5_SIC1_CLB4_CLN1_FKH1_LEU2 | 88 | 0.04% |
| 66 | CLB3_CLB5_SIC1_CLN2_CLB4_LEU2 | 77 | 0.03% |
| 67 | CLB3_CLB5_SIC1_CLN1_CLB3_CLB5_SIC1_CLN1_FKH1_LEU2 | 75 | 0.03% |
| 68 | CLB3_CLB5_SIC1_CLB3_CLB5_LEU2 | 72 | 0.03% |
| 69 | CLB2_FKH2_CLB2_CLN2_FKH2_CLN1_CLB3_CLB5_SIC1_FKH1_LEU2 | 71 | 0.03% |
| 70 | CLN2_CLB2_FKH2_CLB3_CLB5_SIC1_CLB4_LEU2 | 68 | 0.03% |
| 71 | CLB2_CLN1_CLB4_CLB3_CLB5_SIC1_LEU2 | 68 | 0.03% |
| 72 | CLB5_SIC1_CLN1_CLN1_FKH1_LEU2 | 65 | 0.03% |
| 73 | CLB2_FKH2_CLB3_CLB5_SIC1_CLB4_SIC1_CLB4_FKH1_LEU2 | 62 | 0.03% |
| 74 | FKH2_CLB2_CLB3_CLB5_LEU2 | 59 | 0.02% |
| 75 | CLN2_CLB4_CLB3_CLB5_SIC1_CLN1_LEU2 | 55 | 0.02% |
| 76 | CLB5_SIC1_CLB4_FKH1_LEU2 | 53 | 0.02% |
| 77 | CLN2_FKH2_CLB3_CLB5_SIC1_LEU2 | 52 | 0.02% |
| 78 | FKH2_CLB3_CLB5_SIC1_CLN1_LEU2 | 50 | 0.02% |
| 79 | CLN2_CLB3_CLB2_FKH2_CLB3_CLB5_SIC1_CLN1_LEU2 | 45 | 0.02% |
| 80 | FKH2_CLB3_CLB5_SIC1_FKH1_LEU2 | 45 | 0.02% |
| 81 | CLB2_FKH2_CLB3_CLB5_SIC1_CLB4_LEU2 | 42 | 0.02% |
| 82 | CLN2_CLB2_FKH2_LEU2 | 41 | 0.02% |
| 83 | CLB4_CLB3_CLB5_SIC1_CLB3_CLB5_SIC1_CLB4_CLN1_FKH1_LEU2 | 41 | 0.02% |
| 84 | FKH2_CLB3_CLB5_SIC1_CLN1_FKH1_LEU2 | 40 | 0.02% |
| 85 | CLN2_CLN1_FKH1_LEU2 | 39 | 0.02% |
| 86 | CLB2_FKH2_CLN1_LEU2 | 38 | 0.02% |
| 87 | CLN2_CLB3_CLB5_SIC1_CLB2_FKH2_FKH1_LEU2 | 36 | 0.01% |
| 88 | CLB3_CLB5_FKH1_FKH1_LEU2 | 35 | 0.01% |
| 89 | CLB3_CLB5_SIC1_CLB2_FKH2_LEU2 | 35 | 0.01% |
| 90 | CLN2_CLB3_CLB2_FKH2_CLB3_CLB5_SIC1_FKH1_LEU2 | 34 | 0.01% |
| 91 | CLB3_CLB5_SIC1_CLB4_CLN1_FKH1_LEU2 | 33 | 0.01% |
| 92 | CLN2_CLB3_CLB2_FKH2_CLB3_CLB5_SIC1_CLB4_FKH1_LEU2 | 33 | 0.01% |
| 93 | CLB3_CLB5_SIC1_CLB4_LEU2 | 32 | 0.01% |
| 94 | CLB5_SIC1_CLB4_CLN1_LEU2 | 30 | 0.01% |
| 95 | FKH2_CLB3_CLB5_SIC1_CLB4_FKH1_LEU2 | 30 | 0.01% |
| 96 | CLN2_CLB3_CLB5_SIC1_CLB4_LEU2 | 30 | 0.01% |
| 97 | CLB2_FKH2_CLN1_FKH1_LEU2 | 29 | 0.01% |
| 98 | CLB3_CLN2_FKH2_CLB3_CLB5_SIC1_CLN1_LEU2 | 29 | 0.01% |
| 99 | CLN2_CLB3_CLB5_SIC1_CLB4_SIC1_CLB4_FKH1_LEU2 | 28 | 0.01% |
| 100 | CLN2_CLB4_CLB3_CLB5_SIC1_CLN1_FKH1_LEU2 | 28 | 0.01% |
| 101 | CLB2_FKH2_CLB3_CLB5_FKH1_CLB4_LEU2 | 27 | 0.01% |
| 102 | CLB3_CLB5_CLB2_FKH2_CLB2_LEU2 | 25 | 0.01% |
| 103 | CLN2_CLB3_CLB2_FKH2_CLB3_CLB5_SIC1_CLB4_CLN1_LEU2 | 24 | 0.01% |
| 104 | CLN2_CLB2_FKH2_CLB3_CLB5_SIC1_CLB2_FKH2_FKH1_LEU2 | 22 | 0.01% |
| 105 | CLN2_CLB2_CLN2_CLB5_SIC1_LEU2 | 21 | 0.01% |
| 106 | CLB2_FKH2_CLB3_CLB5_SIC1_CLB2_FKH2_FKH1_LEU2 | 21 | 0.01% |
| 107 | CLN2_CLB4_CLB3_CLB5_SIC1_CLB4_FKH1_LEU2 | 21 | 0.01% |
| 108 | CLN1_CLB3_CLB5_SIC1_CLN1_LEU2 | 21 | 0.01% |
| 109 | CLN2_CLB2_FKH2_CLB4_FKH1_LEU2 | 21 | 0.01% |
| 110 | CLN2_CLB2_FKH2_CLB3_CLB5_LEU2 | 20 | 0.01% |
| 111 | CLB2_CLN2_CLB5_SIC1_FKH1_LEU2 | 19 | 0.01% |
| 112 | CLN2_CLN2_CLB3_CLB5_SIC1_FKH1_LEU2 | 18 | 0.01% |
| 113 | CLB3_CLN2_FKH2_CLB3_CLB5_SIC1_FKH1_LEU2 | 18 | 0.01% |
| 114 | CLN2_CLB2_FKH2_CLN1_LEU2 | 17 | 0.01% |
| 115 | CLN2_CLB5_SIC1_CLN1_LEU2 | 17 | 0.01% |
| 116 | CLB2_CLB3_CLB5_SIC1_LEU2 | 17 | 0.01% |
| 117 | CLB2_FKH2_CLB3_CLB5_CLB2_FKH2_LEU2 | 17 | 0.01% |
| 118 | CLB3_CLN2_FKH2_CLB3_CLB5_SIC1_CLB4_FKH1_LEU2 | 17 | 0.01% |
| 119 | CLN2_CLB3_CLB2_FKH2_CLB3_CLB5_SIC1_CLN1_FKH1_LEU2 | 17 | 0.01% |
| 120 | CLN2_CLB2_FKH2_CLB5_LEU2 | 16 | 0.01% |
| 121 | CLN2_CLB3_CLB2_FKH2_FKH1_LEU2 | 16 | 0.01% |
| 122 | FKH1_SIC1_CLB3_CLB5_LEU2 | 15 | 0.01% |
| 123 | CLB2_FKH2_CLB5_SIC1_CLB4_LEU2 | 15 | 0.01% |
| 124 | CLB3_CLB5_SIC1_CLB2_FKH2_FKH1_LEU2 | 14 | 0.01% |
| 125 | CLB2_FKH2_CLN1_CLB3_CLB5_SIC1_FKH1_LEU2 | 14 | 0.01% |
| 126 | CLN2_CLB2_FKH2_CLB3_CLB5_SIC1_CLB3_CLB5_SIC1_CLN1_FKH1_LEU2 | 13 | 0.01% |
| 127 | CLB2_CLB5_SIC1_CLN1_LEU2 | 13 | 0.01% |
| 128 | CLB3_CLB5_CLB5_SIC1_CLB3_CLB5_SIC1_LEU2 | 13 | 0.01% |
| 129 | CLB2_FKH2_CLB3_CLB5_SIC1_CLB3_CLB5_SIC1_CLB4_LEU2 | 13 | 0.01% |
| 130 | CLN2_CLB4_CLB3_CLB5_SIC1_CLB4_CLN1_LEU2 | 13 | 0.01% |
| 131 | CLB3_CLB5_SIC1_CLB3_LEU2 | 13 | 0.01% |
| 132 | CLN2_CLB2_CLN2_CLB5_SIC1_CLB4_FKH1_LEU2 | 12 | 0.00% |
| 133 | CLN2_CLB4_CLN1_FKH1_LEU2 | 12 | 0.00% |
| 134 | FKH2_CLB3_CLB5_LEU2 | 12 | 0.00% |
| 135 | CLB2_FKH2_CLB3_CLB5_FKH1_LEU2 | 12 | 0.00% |
| 136 | CLN2_CLB2_FKH2_CLN1_FKH1_LEU2 | 12 | 0.00% |
| 137 | CLN2_CLB2_CLN2_CLB5_SIC1_CLN1_LEU2 | 12 | 0.00% |
| 138 | FKH2_CLB3_CLB5_SIC1_CLB4_CLN1_LEU2 | 12 | 0.00% |
| 139 | CLB4_CLB2_FKH2_CLB5_LEU2 | 11 | 0.00% |
| 140 | CLN2_CLB4_SIC1_CLB4_FKH1_LEU2 | 11 | 0.00% |
| 141 | FKH2_CLB5_SIC1_CLN1_LEU2 | 10 | 0.00% |
| 142 | CLB4_SIC1_CLB4_FKH1_LEU2 | 9 | 0.00% |
| 143 | CLN2_CLB2_FKH2_CLB3_CLB5_FKH1_LEU2 | 9 | 0.00% |
| 144 | CLB3_CLN2_FKH2_CLB3_CLB5_SIC1_CLN1_FKH1_LEU2 | 9 | 0.00% |
| 145 | CLB2_FKH2_CLB5_SIC1_CLB4_SIC1_CLB4_FKH1_LEU2 | 9 | 0.00% |
| 146 | CLB2_CLB3_CLB5_SIC1_CLN1_LEU2 | 9 | 0.00% |
| 147 | CLB4_CLB3_CLB5_SIC1_LEU2 | 8 | 0.00% |
| 148 | CLB3_CLB5_SIC1_CLB4_SIC1_CLB4_FKH1_LEU2 | 8 | 0.00% |
| 149 | CLN2_CLB5_SIC1_CLB4_CLN1_LEU2 | 8 | 0.00% |
| 150 | CLB3_CLN2_CLB3_CLB5_SIC1_FKH1_LEU2 | 8 | 0.00% |
| 151 | FKH2_CLB3_CLB5_SIC1_CLB3_CLB5_SIC1_FKH1_LEU2 | 8 | 0.00% |
| 152 | CLN1_CLB3_CLB5_SIC1_CLB4_FKH1_LEU2 | 8 | 0.00% |
| 153 | CLB3_CLN2_CLB2_FKH2_CLB3_CLB5_SIC1_LEU2 | 8 | 0.00% |
| 154 | CLN2_CLB5_SIC1_CLB4_FKH1_LEU2 | 7 | 0.00% |
| 155 | CLN2_FKH2_CLN1_CLB3_CLB5_SIC1_FKH1_LEU2 | 7 | 0.00% |
| 156 | CLN2_CLB4_CLB3_CLB5_SIC1_CLB2_FKH2_CLB3_CLB5_SIC1_LEU2 | 7 | 0.00% |
| 157 | CLB2_FKH2_CLB3_CLB5_SIC1_CLB3_CLB5_SIC1_CLN1_FKH1_LEU2 | 7 | 0.00% |
| 158 | CLN2_CLB3_CLB5_SIC1_CLB3_CLB5_SIC1_CLB4_LEU2 | 7 | 0.00% |
| 159 | CLN1_CLB3_CLB5_SIC1_CLB4_CLN1_LEU2 | 7 | 0.00% |
| 160 | CLN2_CLB2_FKH2_CLB3_CLB5_SIC1_CLN2_LEU2 | 7 | 0.00% |
| 161 | SIC1_FKH2_CLB2_FKH2_CLB3_CLB5_SIC1_LEU2 | 7 | 0.00% |
| 162 | CLB2_FKH2_CLB3_CLB2_FKH2_CLB3_CLB5_SIC1_LEU2 | 7 | 0.00% |
| 163 | CLN2_CLB3_CLB5_SIC1_CLB3_CLB5_SIC1_CLN1_FKH1_LEU2 | 7 | 0.00% |
| 164 | CLN2_CLB2_FKH2_CLB3_CLB5_SIC1_CLB4_SIC1_LEU2 | 7 | 0.00% |
| 165 | CLN2_CLB2_FKH2_CLB3_CLB5_SIC1_CLB4_SIC1_FKH1_LEU2 | 6 | 0.00% |
| 166 | CLB2_CLB5_SIC1_CLB4_CLN1_LEU2 | 6 | 0.00% |
| 167 | CLB2_FKH2_CLB3_CLB5_SIC1_CLN2_CLB4_FKH1_LEU2 | 6 | 0.00% |
| 168 | CLN2_CLB2_CLB3_CLB5_SIC1_CLB4_CLN1_LEU2 | 6 | 0.00% |
| 169 | CLB5_SIC1_CLB4_CLN1_FKH1_LEU2 | 6 | 0.00% |
| 170 | CLN2_CLB2_FKH2_CLB3_CLB5_SIC1_CLB4_SIC1_CLB4_LEU2 | 6 | 0.00% |
| 171 | CLB2_CLB5_SIC1_CLB4_FKH1_LEU2 | 6 | 0.00% |
| 172 | CLN2_CLB2_FKH2_CLB5_CLB2_FKH2_CLB2_FKH2_LEU2 | 6 | 0.00% |
| 173 | CLB2_FKH2_CLB5_SIC1_CLB2_FKH2_FKH1_LEU2 | 6 | 0.00% |
| 174 | CLN2_CLB2_FKH2_CLB5_CLN1_LEU2 | 6 | 0.00% |
| 175 | CLB3_CLB2_FKH2_CLB3_CLB5_SIC1_LEU2 | 6 | 0.00% |
| 176 | FKH2_CLB3_CLB5_SIC1_CLB3_CLB5_SIC1_LEU2 | 5 | 0.00% |
| 177 | CLN2_CLB3_CLB5_SIC1_CLN2_CLB4_FKH1_LEU2 | 5 | 0.00% |
| 178 | CLB3_CLN2_FKH2_CLB3_CLB5_SIC1_CLB4_CLN1_LEU2 | 5 | 0.00% |
| 179 | CLN2_CLB5_SIC1_CLN1_FKH1_LEU2 | 5 | 0.00% |
| 180 | CLN2_CLB3_CLB5_SIC1_CLN2_LEU2 | 5 | 0.00% |
| 181 | CLB2_FKH2_CLB3_CLB5_SIC1_CLN2_LEU2 | 5 | 0.00% |
| 182 | FKH2_CLB3_CLB5_SIC1_CLB3_CLB5_SIC1_CLB4_FKH1_LEU2 | 5 | 0.00% |
| 183 | CLN2_CLB2_FKH2_CLB5_SIC1_CLB4_FKH1_LEU2 | 5 | 0.00% |
| 184 | CLB5_SIC1_CLB2_FKH2_FKH1_LEU2 | 5 | 0.00% |
| 185 | CLN2_CLB2_FKH2_CLN1_CLB3_CLB5_SIC1_FKH1_LEU2 | 5 | 0.00% |
| 186 | CLN2_CLB3_CLB2_FKH2_CLB3_CLB5_SIC1_CLB4_SIC1_CLB4_FKH1_LEU2 | 4 | 0.00% |
| 187 | CLN2_CLB3_CLN2_FKH2_CLB3_CLB5_SIC1_LEU2 | 4 | 0.00% |
| 188 | CLN2_CLB2_FKH2_CLB3_CLB5_SIC1_CLB4_SIC1_CLN1_LEU2 | 4 | 0.00% |
| 189 | CLB4_CLB3_CLB5_SIC1_CLN1_LEU2 | 4 | 0.00% |
| 190 | CLN2_CLB2_FKH2_CLB3_CLB5_SIC1_CLN2_CLB4_FKH1_LEU2 | 4 | 0.00% |
| 191 | CLN2_CLB2_CLB3_CLB5_LEU2 | 4 | 0.00% |
| 192 | CLB2_CLB3_CLB5_SIC1_FKH1_LEU2 | 4 | 0.00% |
| 193 | CLN2_CLB2_CLN2_CLB5_SIC1_CLB4_CLN1_FKH1_LEU2 | 4 | 0.00% |
| 194 | CLN2_CLB2_FKH2_CLB3_CLB5_CLB2_FKH2_FKH1_LEU2 | 4 | 0.00% |
| 195 | CLN2_SIC1_CLB4_FKH1_LEU2 | 4 | 0.00% |
| 196 | CLN2_CLB2_CLN2_CLB5_SIC1_CLN1_FKH1_LEU2 | 4 | 0.00% |
| 197 | CLN2_CLB3_CLB2_FKH2_CLB5_SIC1_CLN1_LEU2 | 4 | 0.00% |
| 198 | CLN2_CLB2_FKH2_CLB3_CLB5_SIC1_CLB3_CLB5_SIC1_LEU2 | 4 | 0.00% |
| 199 | CLN2_CLB4_CLB3_CLB5_LEU2 | 4 | 0.00% |
| 200 | CLB3_CLB5_CLB2_FKH2_CLB3_CLB5_SIC1_LEU2 | 4 | 0.00% |
| 201 | CLN2_FKH2_CLB3_CLB5_LEU2 | 4 | 0.00% |
| 202 | CLN2_CLB2_CLB5_SIC1_FKH1_LEU2 | 4 | 0.00% |
| 203 | CLN2_CLB4_CLB3_CLB5_SIC1_CLB2_FKH2_CLB3_CLB5_SIC1_CLN1_LEU2 | 4 | 0.00% |
| 204 | CLN1_CLB3_CLB5_SIC1_CLN1_FKH1_LEU2 | 4 | 0.00% |
| 205 | CLN2_CLB2_FKH2_CLB5_SIC1_LEU2 | 4 | 0.00% |
| 206 | CLN2_CLB3_CLB5_CLB4_FKH1_LEU2 | 3 | 0.00% |
| 207 | CLN2_CLB2_FKH2_CLB5_SIC1_CLN1_FKH1_LEU2 | 3 | 0.00% |
| 208 | CLB4_CLB3_CLB5_SIC1_FKH1_LEU2 | 3 | 0.00% |
| 209 | CLB2_CLB3_CLB5_SIC1_CLB4_FKH1_LEU2 | 3 | 0.00% |
| 210 | CLN2_CLB2_FKH2_CLB3_CLB5_SIC1_CLB3_CLB5_SIC1_FKH1_LEU2 | 3 | 0.00% |
| 211 | CLN2_CLB2_FKH2_CLB3_CLB2_FKH2_CLB3_CLB5_SIC1_LEU2 | 3 | 0.00% |
| 212 | CLN2_CLB4_CLB3_CLB5_SIC1_CLB4_CLN1_FKH1_LEU2 | 3 | 0.00% |
| 213 | CLN2_CLB3_CLB2_FKH2_CLB5_SIC1_CLB4_CLN1_FKH1_LEU2 | 3 | 0.00% |
| 214 | CLB2_FKH2_CLN1_CLB3_CLB5_SIC1_LEU2 | 3 | 0.00% |
| 215 | SIC1_CLB3_CLN1_FKH1_LEU2 | 3 | 0.00% |
| 216 | CLB3_CLB5_SIC1_CLB3_CLB5_SIC1_CLB4_LEU2 | 3 | 0.00% |
| 217 | CLB2_CLB3_CLB5_SIC1_CLN1_FKH1_LEU2 | 3 | 0.00% |
| 218 | CLB2_CLB5_SIC1_CLB4_CLN1_FKH1_LEU2 | 3 | 0.00% |
| 219 | CLN2_CLB5_SIC1_CLB4_CLN1_FKH1_LEU2 | 3 | 0.00% |
| 220 | FKH2_CLB2_FKH2_CLB3_CLB5_SIC1_CLB4_FKH1_LEU2 | 3 | 0.00% |
| 221 | CLB3_CLB5_CLB2_FKH2_CLB3_CLB5_SIC1_CLN1_LEU2 | 3 | 0.00% |
| 222 | CLB3_CLB5_SIC1_CLN1_CLB3_CLB5_SIC1_LEU2 | 3 | 0.00% |
| 223 | CLB2_FKH2_CLB3_CLN1_LEU2 | 3 | 0.00% |
| 224 | CLN2_CLB2_CLN2_CLB3_CLB5_SIC1_FKH1_LEU2 | 3 | 0.00% |
| 225 | FKH2_CLB3_CLB5_SIC1_CLB3_CLB5_SIC1_CLN1_LEU2 | 3 | 0.00% |
| 226 | CLB2_FKH2_CLB4_CLN1_LEU2 | 3 | 0.00% |
| 227 | CLB3_CLN1_CLB5_FKH2_LEU2 | 3 | 0.00% |
| 228 | CLN2_CLB2_FKH2_CLB3_CLB5_CLN1_LEU2 | 3 | 0.00% |
| 229 | CLN2_CLB2_CLN2_CLB5_SIC1_CLB4_CLN1_LEU2 | 3 | 0.00% |
| 230 | CLN2_CLB2_FKH2_CLB3_CLB5_SIC1_CLB2_FKH2_CLB3_CLB5_SIC1_FKH1_LEU2 | 3 | 0.00% |
| 231 | CLB2_CLB5_SIC1_CLN1_FKH1_LEU2 | 3 | 0.00% |
| 232 | CLB2_FKH2_CLB3_CLB5_SIC1_CLB4_SIC1_CLN1_LEU2 | 3 | 0.00% |
| 233 | CLB3_CLN2_FKH2_FKH1_LEU2 | 3 | 0.00% |
| 234 | CLN2_CLB2_CLN1_LEU2 | 2 | 0.00% |
| 235 | CLN2_CLB3_CLB2_FKH2_CLB3_CLB5_SIC1_CLB3_CLB5_SIC1_CLB4_LEU2 | 2 | 0.00% |
| 236 | CLN2_CLB2_FKH2_SIC1_CLB4_FKH1_LEU2 | 2 | 0.00% |
| 237 | CLN2_FKH2_CLB3_CLB5_SIC1_FKH1_LEU2 | 2 | 0.00% |
| 238 | CLB2_FKH2_CLB5_SIC1_CLB3_CLB5_SIC1_CLB4_LEU2 | 2 | 0.00% |
| 239 | CLN2_CLB4_CLB3_CLB5_SIC1_CLB4_SIC1_CLB4_FKH1_LEU2 | 2 | 0.00% |
| 240 | CLB2_FKH2_CLB3_CLB5_CLB4_FKH1_LEU2 | 2 | 0.00% |
| 241 | CLN2_CLB4_CLB3_CLB5_SIC1_CLN2_LEU2 | 2 | 0.00% |
| 242 | CLN2_CLB2_CLN2_LEU2 | 2 | 0.00% |
| 243 | FKH2_CLB2_CLB3_CLB5_SIC1_FKH1_LEU2 | 2 | 0.00% |
| 244 | CLN2_FKH2_CLB3_CLB5_SIC1_CLB4_FKH1_LEU2 | 2 | 0.00% |
| 245 | CLN2_CLB5_SIC1_CLB2_FKH2_FKH1_LEU2 | 2 | 0.00% |
| 246 | CLB2_FKH2_CLB3_LEU2 | 2 | 0.00% |
| 247 | CLN2_CLB3_CLB2_FKH2_CLB3_CLB5_LEU2 | 2 | 0.00% |
| 248 | CLB4_CLB3_CLB5_SIC1_CLB2_FKH2_FKH1_LEU2 | 2 | 0.00% |
| 249 | FKH2_CLN1_CLB3_CLB5_SIC1_CLN1_LEU2 | 2 | 0.00% |
| 250 | CLB2_FKH2_CLB3_CLB5_SIC1_CLN1_CLN1_FKH1_LEU2 | 2 | 0.00% |
| 251 | CLN2_CLB2_FKH2_CLB5_SIC1_FKH1_LEU2 | 2 | 0.00% |
| 252 | CLB2_FKH2_CLB2_LEU2 | 2 | 0.00% |
| 253 | CLN2_CLB2_FKH2_CLB2_CLN2_FKH2_CLN1_CLB3_CLB5_SIC1_FKH1_LEU2 | 2 | 0.00% |
| 254 | CLN2_CLB2_FKH2_CLB3_CLB5_CLB4_CLN2_CLB2_LEU2 | 2 | 0.00% |
| 255 | FKH2_CLB3_CLB5_SIC1_CLB3_CLB5_SIC1_CLB4_CLN1_LEU2 | 2 | 0.00% |
| 256 | CLN2_CLB3_CLB5_SIC1_CLN1_CLB3_CLB5_SIC1_FKH1_LEU2 | 2 | 0.00% |
| 257 | CLB2_FKH2_CLB3_CLB5_SIC1_CLN1_CLB3_CLB5_SIC1_FKH1_LEU2 | 2 | 0.00% |
| 258 | CLN2_CLB2_FKH2_CLB3_CLB5_SIC1_CLB4_SIC1_CLN1_FKH1_LEU2 | 2 | 0.00% |
| 259 | CLN1_CLB3_CLB5_SIC1_CLN1_CLB3_CLB5_SIC1_CLN1_LEU2 | 2 | 0.00% |
| 260 | CLN2_CLB2_FKH2_CLB3_CLB5_SIC1_CLB4_SIC1_CLB4_CLN1_LEU2 | 2 | 0.00% |
| 261 | CLB3_CLN2_FKH2_CLB3_CLB5_SIC1_CLB4_CLN1_FKH1_LEU2 | 2 | 0.00% |
| 262 | CLB4_CLB3_CLB5_SIC1_CLB3_CLB5_SIC1_CLN1_FKH1_LEU2 | 2 | 0.00% |
| 263 | CLB2_FKH2_CLB2_CLN2_FKH2_CLN1_CLB3_CLB5_SIC1_LEU2 | 2 | 0.00% |
| 264 | CLN2_CLB3_CLB2_FKH2_CLB3_CLB5_SIC1_CLB4_CLN1_FKH1_LEU2 | 2 | 0.00% |
| 265 | FKH2_CLB3_CLB5_SIC1_CLB4_SIC1_CLB4_FKH1_LEU2 | 2 | 0.00% |
| 266 | CLB3_CLB5_SIC1_CLN1_CLB3_CLB5_SIC1_FKH1_LEU2 | 2 | 0.00% |
| 267 | CLB3_CLB5_SIC1_CLB3_CLB5_SIC1_CLN1_FKH1_LEU2 | 2 | 0.00% |
| 268 | CLN2_CLB4_CLB3_CLB2_FKH2_CLB3_CLB5_SIC1_LEU2 | 2 | 0.00% |
| 269 | CLB3_CLB5_SIC1_CLN1_CLB3_CLB5_SIC1_CLB4_FKH1_LEU2 | 2 | 0.00% |
| 270 | CLN2_CLB3_CLB5_CLB2_FKH2_FKH1_LEU2 | 2 | 0.00% |
| 271 | CLN2_CLB5_SIC1_CLB4_LEU2 | 2 | 0.00% |
| 272 | CLB3_CLB5_SIC1_CLN1_CLB3_CLB5_SIC1_CLN1_LEU2 | 2 | 0.00% |
| 273 | CLB3_CLN2_CLB2_FKH2_CLB3_CLB5_SIC1_CLB4_CLN1_LEU2 | 2 | 0.00% |
| 274 | CLN1_CLB3_CLB5_SIC1_CLB4_LEU2 | 2 | 0.00% |
| 275 | CLB2_FKH2_CLB4_CLN1_FKH1_LEU2 | 2 | 0.00% |
| 276 | CLN2_CLB2_FKH2_CLB4_LEU2 | 2 | 0.00% |
| 277 | CLB4_CLB3_CLB5_SIC1_CLB4_CLN1_LEU2 | 2 | 0.00% |
| 278 | CLN2_CLB2_CLB4_CLN1_LEU2 | 2 | 0.00% |
| 279 | CLB5_SIC1_CLN1_LEU2 | 2 | 0.00% |
| 280 | CLB3_CLB5_CLB2_FKH2_CLB3_CLB5_SIC1_CLB4_FKH1_LEU2 | 1 | 0.00% |
| 281 | CLN2_CLB2_FKH2_CLB3_CLN1_CLB5_FKH2_LEU2 | 1 | 0.00% |
| 282 | CLB2_FKH2_CLB2_CLN2_FKH2_CLN1_CLB3_CLB5_SIC1_CLB4_FKH1_LEU2 | 1 | 0.00% |
| 283 | CLN2_CLB2_SIC1_SIC1_CLB4_FKH1_LEU2 | 1 | 0.00% |
| 284 | CLB2_FKH2_CLB3_SIC1_LEU2 | 1 | 0.00% |
| 285 | CLN2_CLB2_FKH2_CLB3_CLB5_SIC1_CLB4_CLB4_FKH1_LEU2 | 1 | 0.00% |
| 286 | CLN2_CLB2_FKH2_CLB3_CLB5_SIC1_CLB3_CLB5_SIC1_CLB4_CLN1_FKH1_LEU2 | 1 | 0.00% |
| 287 | CLN2_CLB2_CLN2_FKH2_CLN1_CLB3_CLB5_SIC1_FKH1_LEU2 | 1 | 0.00% |
| 288 | CLN2_CLB4_CLB3_CLB5_SIC1_CLB4_LEU2 | 1 | 0.00% |
| 289 | CLB2_FKH2_CLB3_CLB5_SIC1_CLB3_CLB5_SIC1_FKH1_LEU2 | 1 | 0.00% |
| 290 | CLN2_CLB3_CLB5_SIC1_CLB4_FKH1_LEU2_CLB2_CLB3_CLB5_SIC1_CLB4_CLN1_LEU2 | 1 | 0.00% |
| 291 | CLB2_FKH2_CLB3_FKH1_LEU2 | 1 | 0.00% |
| 292 | CLB5_SIC1_CLB3_CLB2_FKH2_LEU2 | 1 | 0.00% |
| 293 | FKH2_CLB3_CLB5_SIC1_CLN2_LEU2 | 1 | 0.00% |
| 294 | CLN2_CLB3_CLB2_FKH2_CLB3_CLB5_SIC1_CLB3_CLB5_SIC1_CLN1_FKH1_LEU2 | 1 | 0.00% |
| 295 | CLB2_FKH2_CLB3_CLB5_CLN1_LEU2 | 1 | 0.00% |
| 296 | CLN2_CLB4_CLB3_CLB5_SIC1_CLB2_FKH2_CLB3_CLB5_SIC1_CLB4_CLN1_LEU2 | 1 | 0.00% |
| 297 | FKH2_CLB3_CLB5_FKH1_LEU2 | 1 | 0.00% |
| 298 | CLB2_CLB3_CLB5_SIC1_CLB3_CLB5_SIC1_FKH1_LEU2 | 1 | 0.00% |
| 299 | CLN2_CLB3_CLB5_SIC1_CLB3_CLB5_SIC1_LEU2 | 1 | 0.00% |
| 300 | CLB2_FKH2_CLB2_CLN2_CLB2_FKH2_CLB3_CLB5_SIC1_LEU2 | 1 | 0.00% |
| 301 | CLB3_CLB5_CLB2_FKH2_CLB3_CLB5_SIC1_CLN1_FKH1_LEU2 | 1 | 0.00% |
| 302 | CLN2_CLB2_FKH2_CLB3_CLN2_FKH2_CLB3_CLB5_SIC1_LEU2 | 1 | 0.00% |
| 303 | CLB3_CLN2_CLB2_FKH2_CLB3_CLB5_SIC1_CLB4_FKH1_LEU2 | 1 | 0.00% |
| 304 | CLB2_FKH2_CLB4_FKH1_LEU2 | 1 | 0.00% |
| 305 | CLB2_FKH2_CLB2_FKH2_CLB3_CLB5_SIC1_CLN1_LEU2 | 1 | 0.00% |
| 306 | CLB2_FKH2_CLB3_CLB5_SIC1_CLN1_LEU2_LEU2 | 1 | 0.00% |
| 307 | CLN2_CLB3_CLB5_CLN1_LEU2 | 1 | 0.00% |
| 308 | SIC1_CLB2_FKH2_FKH1_LEU2 | 1 | 0.00% |
| 309 | CLB3_CLN2_CLB4_CLB3_CLB5_SIC1_CLB2_FKH2_FKH1_LEU2 | 1 | 0.00% |
| 310 | CLB2_CLB5_SIC1_FKH1_LEU2_CLB3_CLB5_SIC1_CLN1_FKH1_LEU2 | 1 | 0.00% |
| 311 | CLN2_CLB2_FKH2_CLB3_CLB5_SIC1_LEU2_CLB5_SIC1_CLN1_LEU2 | 1 | 0.00% |
| 312 | CLB2_FKH2_CLB3_CLN2_FKH2_CLB3_CLB5_SIC1_LEU2 | 1 | 0.00% |
| 313 | CLB3_CLN2_CLB3_CLB5_SIC1_CLB4_FKH1_LEU2 | 1 | 0.00% |
| 314 | CLB3_CLB5_SIC1_CLB2_FKH2_CLB3_CLB5_SIC1_FKH1_LEU2 | 1 | 0.00% |
| 315 | CLN2_CLB2_CLN2_CLB2_FKH2_CLB3_CLB5_SIC1_LEU2 | 1 | 0.00% |
| 316 | CLN2_CLN1_CLB3_CLB5_SIC1_LEU2 | 1 | 0.00% |
| 317 | CLN2_CLB4_CLB3_CLB5_SIC1_CLB4_SIC1_FKH1_LEU2 | 1 | 0.00% |
| 318 | CLB3_CLN2_FKH2_CLB3_CLB5_SIC1_CLB4_SIC1_CLB4_FKH1_LEU2 | 1 | 0.00% |
| 319 | CLN2_SIC1_CLB3_CLB5_SIC1_LEU2 | 1 | 0.00% |
| 320 | CLN2_CLB3_CLB2_CLN2_CLB5_SIC1_FKH1_LEU2 | 1 | 0.00% |
| 321 | CLB4_CLB3_CLB5_SIC1_CLB3_CLB5_SIC1_FKH1_LEU2 | 1 | 0.00% |
| 322 | CLB3_CLB5_SIC1_CLB3_CLB5_SIC1_CLB4_FKH1_LEU2 | 1 | 0.00% |
| 323 | CLB3_CLN2_FKH2_CLB5_SIC1_CLN1_LEU2 | 1 | 0.00% |
| 324 | CLN2_CLB4_CLB3_CLB5_SIC1_CLB2_FKH2_CLB3_CLB5_SIC1_CLB4_CLN1_FKH1_LEU2 | 1 | 0.00% |
| 325 | CLN2_CLB3_CLB2_FKH2_CLB5_CLN1_LEU2 | 1 | 0.00% |
| 326 | CLB4_CLB3_CLB5_SIC1_CLB3_CLB5_SIC1_CLB4_CLN1_LEU2 | 1 | 0.00% |
| 327 | CLB2_CLB3_CLB5_SIC1_CLB4_CLN1_FKH1_LEU2 | 1 | 0.00% |
| 328 | CLN1_FKH1_CLB3_CLB5_SIC1_CLN1_FKH1_LEU2 | 1 | 0.00% |
| 329 | CLN1_CLB3_CLB5_SIC1_CLB4_CLN1_FKH1_LEU2 | 1 | 0.00% |
| 330 | SIC1_CLB3_CLN1_CLB5_SIC1_CLB4_CLN1_LEU2 | 1 | 0.00% |
| 331 | CLN1_CLB3_CLB5_SIC1_CLN2_CLB4_FKH1_LEU2 | 1 | 0.00% |
| 332 | CLN2_CLB3_CLB5_SIC1_FKH1_LEU2_CLN1_LEU2 | 1 | 0.00% |
| 333 | CLB2_FKH2_CLB3_CLB5_SIC1_CLB3_CLB5_LEU2 | 1 | 0.00% |
| 334 | CLN2_CLB2_CLB3_CLB5_SIC1_CLB4_FKH1_LEU2 | 1 | 0.00% |
| 335 | CLB2_FKH2_CLB2_CLN2_CLB3_CLB5_SIC1_CLB4_FKH1_LEU2 | 1 | 0.00% |
| 336 | CLB3_CLN2_CLB5_SIC1_FKH1_LEU2 | 1 | 0.00% |
| 337 | CLB2_FKH2_CLB2_CLN2_FKH2_CLB3_CLB5_SIC1_FKH1_LEU2 | 1 | 0.00% |
| 338 | CLN2_CLB3_CLB5_FKH2_LEU2 | 1 | 0.00% |
| 339 | CLN2_CLB2_FKH2_CLB3_CLB5_FKH1_CLB4_LEU2 | 1 | 0.00% |
| 340 | CLN2_CLB2_FKH2_CLB2_FKH2_CLB3_CLB5_SIC1_CLB4_LEU2 | 1 | 0.00% |
| 341 | FKH2_CLB3_CLB5_SIC1_CLB4_LEU2 | 1 | 0.00% |
| 342 | CLB2_CLN2_FKH2_CLN1_CLB3_CLB5_SIC1_FKH1_LEU2 | 1 | 0.00% |
| 343 | CLN2_CLB3_CLB5_SIC1_CLN1_CLB3_CLB5_SIC1_CLN1_FKH1_LEU2 | 1 | 0.00% |
| 344 | CLB3_CLB5_SIC1_CLN1_CLB3_CLB5_SIC1_CLB4_CLN1_LEU2 | 1 | 0.00% |
| 345 | CLB2_FKH2_CLB2_CLN2_FKH2_CLN1_CLB3_CLB5_SIC1_CLB4_CLN1_FKH1_LEU2 | 1 | 0.00% |
| 346 | CLN2_CLB2_FKH2_CLB2_LEU2 | 1 | 0.00% |
| 347 | CLN2_CLB3_CLB5_SIC1_CLB3_CLB5_FKH1_LEU2 | 1 | 0.00% |
| 348 | CLN2_CLB3_CLB2_FKH2_LEU2 | 1 | 0.00% |
| 349 | CLN2_CLB2_SIC1_CLB4_FKH1_LEU2 | 1 | 0.00% |
| 350 | CLN2_FKH2_CLB3_CLB5_SIC1_CLN1_LEU2 | 1 | 0.00% |
| 351 | LEU2_CLN2_CLB2_FKH2_CLB3_CLB5_SIC1_LEU2 | 1 | 0.00% |
| 352 | CLN2_CLB2_FKH2_CLB3_CLB5_SIC1_CLB3_CLB5_LEU2 | 1 | 0.00% |
| 353 | CLB3_CLN2_CLB2_FKH2_CLB3_CLB5_SIC1_FKH1_LEU2 | 1 | 0.00% |
| 354 | CLB3_CLB2_FKH2_CLB3_CLB5_SIC1_CLN1_FKH1_LEU2 | 1 | 0.00% |
| 355 | CLB4_CLB3_CLB5_SIC1_CLB3_CLB5_SIC1_LEU2 | 1 | 0.00% |
| 356 | CLN1_CLN1_FKH1_LEU2 | 1 | 0.00% |
| 357 | CLN2_CLB2_FKH2_CLB3_CLN1_LEU2 | 1 | 0.00% |
| 358 | CLN2_CLB4_CLN1_LEU2 | 1 | 0.00% |
| 359 | FKH2_CLB5_SIC1_CLB4_CLN1_FKH1_LEU2 | 1 | 0.00% |
| 360 | CLN2_CLB3_CLB5_SIC1_CLB3_CLB5_SIC1_CLB4_FKH1_LEU2 | 1 | 0.00% |
| 361 | CLN2_CLB2_FKH2_CLB3_CLB5_SIC1_CLB3_CLB5_SIC1_CLB4_CLN1_LEU2 | 1 | 0.00% |
| 362 | CLN2_CLB2_FKH2_CLB3_CLB5_CLB2_FKH2_CLB2_LEU2 | 1 | 0.00% |
| 363 | CLB3_CLB5_CLN1_FKH1_LEU2 | 1 | 0.00% |
| 364 | CLB3_CLB5_SIC1_CLN1_CLN1_FKH1_LEU2 | 1 | 0.00% |
| 365 | CLN2_CLB2_FKH2_CLB3_CLB5_SIC1_CLB4_SIC1_CLB4_CLN1_FKH1_LEU2 | 1 | 0.00% |
| 366 | CLB2_FKH2_CLB3_CLB5_SIC1_CLN1_LEU2_CLN2_CLB2_FKH2_CLB3_CLB5_SIC1_FKH1_LEU2 | 1 | 0.00% |
| 367 | CLB2_FKH2_CLB5_SIC1_CLB3_CLB5_LEU2 | 1 | 0.00% |
| 368 | CLB3_CLN2_FKH2_CLB3_CLB5_SIC1_CLB3_CLB5_SIC1_CLN1_FKH1_LEU2 | 1 | 0.00% |
| 369 | CLB2_FKH2_CLB2_FKH2_CLB3_CLB5_SIC1_LEU2 | 1 | 0.00% |
| 370 | CLB2_FKH2_CLB3_CLB5_SIC1_CLB2_FKH2_LEU2 | 1 | 0.00% |
| 371 | CLN2_CLB5_CLB2_FKH2_FKH1_LEU2 | 1 | 0.00% |
| 372 | CLB2_CLB5_SIC1_CLB4_SIC1_CLB4_FKH1_LEU2 | 1 | 0.00% |
| 373 | CLN2_CLB2_CLB5_SIC1_CLB4_FKH1_LEU2 | 1 | 0.00% |
| 374 | CLB3_CLN2_FKH2_LEU2 | 1 | 0.00% |
| 375 | CLN2_CLB3_CLB5_SIC1_CLN2_CLB4_LEU2 | 1 | 0.00% |
| 376 | SIC1_CLB4_CLN1_FKH1_LEU2 | 1 | 0.00% |
| 377 | SIC1_CLN2_CLB4_FKH1_LEU2 | 1 | 0.00% |
| 378 | CLN2_CLB3_CLB5_CLB2_FKH2_LEU2 | 1 | 0.00% |
| 379 | CLB5_SIC1_CLN1_LEU2_LEU2 | 1 | 0.00% |
| 380 | CLN2_CLB3_CLN1_CLB5_FKH2_LEU2 | 1 | 0.00% |
| 381 | FKH2_CLN1_FKH1_LEU2 | 1 | 0.00% |
| 382 | CLN1_CLB5_SIC1_FKH1_LEU2 | 1 | 0.00% |
| 383 | CLN2_CLB4_CLB3_FKH1_LEU2 | 1 | 0.00% |
| 384 | CLN1_CLN1_CLB3_CLB5_SIC1_FKH1_LEU2 | 1 | 0.00% |
| 385 | CLB3_CLB5_SIC1_CLN2_CLB4_CLN1_LEU2 | 1 | 0.00% |
| 386 | CLB3_CLN2_CLB4_FKH1_LEU2 | 1 | 0.00% |
| 387 | CLB2_CLN2_FKH2_CLB5_CLN1_LEU2 | 1 | 0.00% |
| 388 | CLN2_CLB2_FKH2_CLB3_FKH1_LEU2 | 1 | 0.00% |
| 389 | CLN2_CLB3_CLB5_CLB4_LEU2 | 1 | 0.00% |
| 390 | CLN1_FKH1_CLB3_CLB5_SIC1_CLN1_LEU2 | 1 | 0.00% |
| 391 | SIC1_CLB3_CLB5_SIC1_LEU2 | 1 | 0.00% |
| 392 | SIC1_CLB3_CLN1_CLB5_LEU2 | 1 | 0.00% |
| 393 | FKH2_CLB3_CLN1_CLB5_FKH2_LEU2 | 1 | 0.00% |
| 394 | CLN2_CLB2_CLN2_CLB3_CLB5_SIC1_CLB4_FKH1_LEU2 | 1 | 0.00% |
| 395 | SIC1_CLB3_CLB5_SIC1_CLB4_CLN1_LEU2 | 1 | 0.00% |
| 396 | CLB2_CLN2_CLB5_SIC1_LEU2 | 1 | 0.00% |
| 397 | CLN2_CLB2_SIC1_FKH1_LEU2 | 1 | 0.00% |
| 398 | CLN1_FKH1_CLB3_CLB5_SIC1_CLB4_FKH1_LEU2 | 1 | 0.00% |
| 399 | FKH2_CLN1_CLB3_CLB5_SIC1_CLN1_FKH1_LEU2 | 1 | 0.00% |
| 400 | CLN2_CLB2_FKH2_CLB5_FKH1_LEU2 | 1 | 0.00% |
| 401 | CLB4_CLB3_CLB5_SIC1_CLN1_FKH1_LEU2 | 1 | 0.00% |
| 402 | CLN2_CLB4_CLB3_CLB5_SIC1_CLN2_CLB4_FKH1_LEU2 | 1 | 0.00% |
| 403 | CLB5_CLB4_SIC1_CLB4_FKH1_LEU2 | 1 | 0.00% |
| 404 | CLN1_CLB3_CLB5_SIC1_CLN1_CLB3_CLB5_SIC1_LEU2 | 1 | 0.00% |
| 405 | CLN2_CLB2_FKH2_CLB3_CLB5_SIC1_CLB3_CLB5_SIC1_CLN1_LEU2 | 1 | 0.00% |
| 406 | CLB2_CLB5_CLB2_FKH2_FKH1_LEU2 | 1 | 0.00% |
| 407 | CLN2_CLB2_FKH2_CLB5_SIC1_CLB4_CLN1_LEU2 | 1 | 0.00% |
| 408 | CLN2_CLB2_FKH2_CLB3_CLB5_CLB2_FKH2_LEU2 | 1 | 0.00% |
| 409 | CLN2_CLB2_FKH2_SIC1_FKH1_LEU2 | 1 | 0.00% |
| 410 | CLB3_CLN2_FKH2_CLB3_CLB5_SIC1_CLN2_CLB4_FKH1_LEU2 | 1 | 0.00% |
| 411 | CLN1_FKH1_CLB3_CLB5_SIC1_CLB4_LEU2 | 1 | 0.00% |
| 412 | CLB2_FKH2_CLB5_SIC1_CLN1_CLN1_FKH1_LEU2 | 1 | 0.00% |
| 413 | CLN2_CLB2_FKH2_CLB5_CLB4_FKH1_LEU2 | 1 | 0.00% |
| 414 | CLN2_CLB3_CLB5_SIC1_SIC1_CLB4_FKH1_LEU2 | 1 | 0.00% |
| 415 | FKH2_CLN1_CLB3_CLB5_SIC1_FKH1_LEU2 | 1 | 0.00% |
| 416 | CLB2_FKH2_CLB3_CLB5_SIC1_CLB3_CLB5_SIC1_CLB4_CLN1_FKH1_LEU2 | 1 | 0.00% |
| 417 | CLB3_CLN2_FKH2_CLB3_CLB5_LEU2 | 1 | 0.00% |
| 418 | SIC1_CLB3_CLB5_SIC1_CLN1_LEU2 | 1 | 0.00% |
| 419 | SIC1_CLB3_CLB5_SIC1_CLN1_FKH1_LEU2 | 1 | 0.00% |
| 420 | CLN2_CLB2_FKH2_CLB3_CLB5_CLN1_FKH1_LEU2 | 1 | 0.00% |
| 421 | CLN2_CLB4_CLB3_CLB5_SIC1_CLB2_FKH2_CLB3_CLB5_SIC1_FKH1_LEU2 | 1 | 0.00% |
| 422 | CLB2_CLN2_FKH2_CLN1_CLB3_CLB5_SIC1_LEU2 | 1 | 0.00% |
| 423 | CLN2_CLB2_CLB4_FKH1_LEU2 | 1 | 0.00% |
| 424 | CLB2_FKH2_CLB3_CLB5_SIC1_CLB3_LEU2 | 1 | 0.00% |
| 425 | CLB5_CLB4_CLN2_CLB2_FKH2_CLB3_CLB5_SIC1_LEU2 | 1 | 0.00% |
| 426 | CLN2_CLB2_FKH2_CLN1_CLB3_CLB5_SIC1_LEU2 | 1 | 0.00% |
| 427 | CLN2_CLB2_FKH2_CLB3_CLB5_SIC1_CLB3_CLB5_SIC1_CLB4_FKH1_LEU2 | 1 | 0.00% |
| 428 | FKH2_CLB3_CLB5_SIC1_CLB2_FKH2_FKH1_LEU2 | 1 | 0.00% |
| 429 | CLB4_CLB3_CLB5_SIC1_CLB4_CLN1_FKH1_LEU2 | 1 | 0.00% |
| 430 | CLB2_FKH2_CLB3_CLB5_SIC1_CLN1_CLB3_CLB5_SIC1_CLN1_FKH1_LEU2 | 1 | 0.00% |
| 431 | CLB3_CLB5_SIC1_CLN2_CLN2_CLB4_FKH1_LEU2 | 1 | 0.00% |

**Table S2:** Gene deletion patterns in 25 minimal viable cell cycle strains. Gene deletion patterns in 25 minimal viable cell cycle strains derived from SCRaMbLE-mediated gene deletion(s) in the synthetic 9-gene cluster strain. Strains were identified via multiplex PCR screening and genotyped using long PCR analysis and POLAR-seq. Rows list strains; columns show the 9 target cell cycle genes (*CLN1*, *CLN2*, *FKH1*, *FKH2*, *SIC1*, *CLB2*, *CLB3*, *CLB4*, *CLB5*). X: gene deleted; blank cell indicates gene retained. Peach shading shows strain pairs whose cluster genotypes were revealed to be identical, but the strains were independently isolated and cannot be guaranteed to be identical elsewhere in the genome without having full genome sequence information.

| Genotypes | CLN1 | CLN2 | FKH1 | FKH2 | SIC1 | CLB2 | CLB3 | CLB4 | CLB5 |
| --- | --- | --- | --- | --- | --- | --- | --- | --- | --- |
| yAM032 | X |  |  |  |  |  |  |  |  |
| yAM037 |  | X |  |  |  |  |  |  |  |
| yAM038 |  | X |  |  |  | X |  |  |  |
| yAM042<br>yAM049 |  | X |  |  |  |  |  | X |  |
| yAM063 |  |  | X |  |  |  | X |  |  |
| yAM051<br>yAM064 | X |  |  | X |  | X |  |  |  |
| yAM061 |  | X |  | X |  | X |  |  |  |
| yAM046 |  | X |  |  |  |  | X | X |  |
| yAM098 |  |  | X |  |  |  | X | X |  |
| yAM047<br>yAM099 | X |  | X |  |  |  |  | X |  |
| yAM100 |  | X | X |  |  |  |  | X |  |
| yAM065 | X | X |  |  | X |  |  |  |  |
| yAM035 | X | X |  |  | X |  |  | X |  |
| yAM050 |  | X |  | X |  | X |  | X |  |
| yAM058 | X |  | X |  | X |  |  | X |  |
| yAM048 |  | X | X |  |  | X |  | X |  |
| yAM101 |  | X | X |  |  |  | X | X |  |
| yAM073 | X |  | X | X | X |  |  | X |  |
| yAM074 | X | X | X |  | X |  |  | X |  |
| yAM057<br>yAM034 | X | X | X |  |  |  | X | X |  |
| yAM041 |  | X | X | X |  | X | X | X |  |

**Table S3:** Complete genotypes of 25 minimal viable cell cycle strains. Complete genotypes of 25 minimal viable cell cycle strains listed in Table S2. All strains were derived from the parental BY4741 strain (*MATa, his3*Δ*1, leu2*Δ*0, met15*Δ*0, ura3*Δ*0*) and were verified by multiplex PCR, long PCR, and POLAR-seq.

| Minimal strains verified by Multiplex PCR, long-read PCR and POLAR-seq |  |
| --- | --- |
| yAM032 | *MATa his3Δ1 leu2Δ0 met15Δ0 ura3Δ0 cln2Δ clb2Δ fkh2Δ clb5Δ clb3Δ sic1Δ clb4Δ cln1Δ fkh1Δ CLN2 CLB2 FKH2 CLB5 CLB3 SIC1 CLB4 FKH1 at URA3 locus (URA3)* |
| yAM037 | *MATa his3Δ1 leu2Δ0 met15Δ0 ura3Δ0 cln2Δ clb2Δ fkh2Δ clb5Δ clb3Δ sic1Δ clb4Δ cln1Δ fkh1Δ CLB2 FKH2 CLB5 CLB3 SIC1 CLB4 CLN1 FKH1 at URA3 locus (URA3)* |
| yAM038 | *MATa his3Δ1 leu2Δ0 met15Δ0 ura3Δ0 cln2Δ clb2Δ fkh2Δ clb5Δ clb3Δ sic1Δ clb4Δ cln1Δ fkh1Δ CLN2 FKH2 CLB5 CLB3 SIC1 CLB4 FKH1 at URA3 locus (URA3)* |
| yAM042 | *MATa his3Δ1 leu2Δ0 met15Δ0 ura3Δ0 cln2Δ clb2Δ fkh2Δ clb5Δ clb3Δ sic1Δ clb4Δ cln1Δ fkh1Δ CLB2 FKH2 CLB5 CLB3 SIC1 CLN1 FKH1 at URA3 locus (URA3)* |
| yAM049 | *MATa his3Δ1 leu2Δ0 met15Δ0 ura3Δ0 cln2Δ clb2Δ fkh2Δ clb5Δ clb3Δ sic1Δ clb4Δ cln1Δ fkh1Δ CLB2 FKH2 CLB5 CLB3 SIC1 CLN1 FKH1 at URA3 locus (URA3)* |
| yAM063 | *MATa his3Δ1 leu2Δ0 met15Δ0 ura3Δ0 cln2Δ clb2Δ fkh2Δ clb5Δ clb3Δ sic1Δ clb4Δ cln1Δ fkh1Δ CLN2 CLB2 FKH2 CLB5 SIC1 CLB4 CLN1 at URA3 locus (URA3)* |
| yAM051 | *MATa his3Δ1 leu2Δ0 met15Δ0 ura3Δ0 cln2Δ clb2Δ fkh2Δ clb5Δ clb3Δ sic1Δ clb4Δ cln1Δ fkh1Δ CLN2 CLB5 CLB3 SIC1 CLB4 FKH1 at URA3 locus (URA3)* |
| yAM064 | *MATa his3Δ1 leu2Δ0 met15Δ0 ura3Δ0 cln2Δ clb2Δ fkh2Δ clb5Δ clb3Δ sic1Δ clb4Δ cln1Δ fkh1Δ CLN2 CLB5 CLB3 SIC1 CLB4 FKH1 at URA3 locus (URA3)* |
| yAM061 | *MATa his3Δ1 leu2Δ0 met15Δ0 ura3Δ0 cln2Δ clb2Δ fkh2Δ clb5Δ clb3Δ sic1Δ clb4Δ cln1Δ fkh1Δ CLB5 CLB3 SIC1 CLB4 CLN1 FKH1 at URA3 locus (URA3)* |
| yAM046 | *MATa his3Δ1 leu2Δ0 met15Δ0 ura3Δ0 cln2Δ clb2Δ fkh2Δ clb5Δ clb3Δ sic1Δ clb4Δ cln1Δ fkh1Δ CLB2 FKH2 CLB5 SIC1 CLN1 FKH1 at URA3 locus (URA3)* |
| yAM098 | *MATa his3Δ1 leu2Δ0 met15Δ0 ura3Δ0 cln2Δ clb2Δ fkh2Δ clb5Δ clb3Δ sic1Δ clb4Δ cln1Δ fkh1Δ CLN2 CLB2 FKH2 CLB5 SIC1 CLN1 at URA3 locus (URA3)* |
| yAM047 | *MATa his3Δ1 leu2Δ0 met15Δ0 ura3Δ0 cln2Δ clb2Δ fkh2Δ clb5Δ clb3Δ sic1Δ clb4Δ cln1Δ fkh1Δ CLN2 CLB2 FKH2 CLB5 CLB3 SIC1 at URA3 locus (URA3)* |
| yAM099 | *MATa his3Δ1 leu2Δ0 met15Δ0 ura3Δ0 cln2Δ clb2Δ fkh2Δ clb5Δ clb3Δ sic1Δ clb4Δ cln1Δ fkh1Δ CLN2 CLB2 FKH2 CLB5 CLB3 SIC1 at URA3 locus (URA3)* |
| yAM100 | *MATa his3Δ1 leu2Δ0 met15Δ0 ura3Δ0 cln2Δ clb2Δ fkh2Δ clb5Δ clb3Δ sic1Δ clb4Δ cln1Δ fkh1Δ CLB2 FKH2 CLB5 CLB3 SIC1 CLN1 at URA3 locus (URA3)* |
| yAM065 | *MATa his3Δ1 leu2Δ0 met15Δ0 ura3Δ0 cln2Δ clb2Δ fkh2Δ clb5Δ clb3Δ sic1Δ clb4Δ cln1Δ fkh1Δ CLB2 FKH2 CLB5 CLB3 CLB4 FKH1 at URA3 locus (URA3)* |
| yAM035 | *MATa his3Δ1 leu2Δ0 met15Δ0 ura3Δ0 cln2Δ clb2Δ fkh2Δ clb5Δ clb3Δ sic1Δ clb4Δ cln1Δ fkh1Δ CLB2 FKH2 CLB5 CLB3 FKH1 at URA3 locus (URA3)* |
| yAM050 | *MATa his3Δ1 leu2Δ0 met15Δ0 ura3Δ0 cln2Δ clb2Δ fkh2Δ clb5Δ clb3Δ sic1Δ clb4Δ cln1Δ fkh1Δ CLB5 CLB3 SIC1 CLN1 FKH1 at URA3 locus (URA3)* |
| yAM058 | *MATa his3Δ1 leu2Δ0 met15Δ0 ura3Δ0 cln2Δ clb2Δ fkh2Δ clb5Δ clb3Δ sic1Δ clb4Δ cln1Δ fkh1Δ CLN2 CLB2 FKH2 CLB5 CLB3 at URA3 locus (URA3)* |
| yAM048 | *MATa his3Δ1 leu2Δ0 met15Δ0 ura3Δ0 cln2Δ clb2Δ fkh2Δ clb5Δ clb3Δ sic1Δ clb4Δ cln1Δ fkh1Δ FKH2 CLB5 CLB3 SIC1 CLN1 at URA3 locus (URA3)* |
| yAM101 | *MATa his3Δ1 leu2Δ0 met15Δ0 ura3Δ0 cln2Δ clb2Δ fkh2Δ clb5Δ clb3Δ sic1Δ clb4Δ cln1Δ fkh1Δ CLB2 FKH2 CLB5 SIC1 CLN1 at URA3 locus (URA3)* |
| yAM073 | *MATa his3Δ1 leu2Δ0 met15Δ0 ura3Δ0 cln2Δ clb2Δ fkh2Δ clb5Δ clb3Δ sic1Δ clb4Δ cln1Δ fkh1Δ CLN2 CLB2 CLB5 CLB3 at URA3 locus (URA3)* |
| yAM074 | *MATa his3Δ1 leu2Δ0 met15Δ0 ura3Δ0 cln2Δ clb2Δ fkh2Δ clb5Δ clb3Δ sic1Δ clb4Δ cln1Δ fkh1Δ CLB2 FKH2 CLB5 CLB3 at URA3 locus (URA3)* |
| yAM057 | *MATa his3Δ1 leu2Δ0 met15Δ0 ura3Δ0 cln2Δ clb2Δ fkh2Δ clb5Δ clb3Δ sic1Δ clb4Δ cln1Δ fkh1Δ CLB2 FKH2 CLB5 SIC1 at URA3 locus (URA3)* |
| yAM034 | *MATa his3Δ1 leu2Δ0 met15Δ0 ura3Δ0 cln2Δ clb2Δ fkh2Δ clb5Δ clb3Δ sic1Δ clb4Δ cln1Δ fkh1Δ CLB2 FKH2 CLB5 SIC1 at URA3 locus (URA3)* |
| yAM041 | *MATa his3Δ1 leu2Δ0 met15Δ0 ura3Δ0 cln2Δ clb2Δ fkh2Δ clb5Δ clb3Δ sic1Δ clb4Δ cln1Δ fkh1Δ CLB5 SIC1 CLN1 at URA3 locus (URA3)* |

## Supplementary Note: Stepwise Construction of Synthetic Cell Cycle Gene Clusters

The construction of synthetic gene clusters provides a systematic approach for studying essential cellular functions by relocating functionally related genes together into defined genomic locations. We developed an iterative methodology combining CRISPR/Cas9-mediated gene deletions with homologous recombination-based assembly to construct synthetic cell cycle gene clusters in *Saccharomyces cerevisiae* (**Fig. S1**). Through this approach, we successfully built consecutive clusters containing 3, 5, 6, and 9 cell cycle genes (**Fig. S2**). This supplementary note describes both our detailed experimental methodology and the technical challenges encountered during synthetic cell cycle gene cluster construction.

### Target Gene Selection and Strain Background

The synthetic cluster was designed to include 11 cell cycle genes representing the core regulatory network controlling cyclin waves in budding yeast. Target genes included the G_1_ cyclins *CLN1* and *CLN2*, the complete set of B-type cyclins (*CLB1* through *CLB6*), and key transcriptional regulators *FKH1*, *FKH2*, and the cyclin inhibitor *SIC1*. All strains were constructed using the BY4741 yeast strain (*MATa his3Δ1 leu2Δ0 met15Δ0 ura3Δ0*), chosen for its compatibility with established genome engineering protocols and extensive genomic annotation^1–3^

### CRISPR/Cas9 Deletion Strategy

Native genes were deleted using a precise CRISPR/Cas9-mediated approach involving two guide RNAs targeting sequences immediately upstream and downstream of each gene’s open reading frame. The CRISPR/Cas9 system, delivered via plasmid transformation, introduces double-strand breaks at the 5’ and 3’ ends of the target gene. The deleted ORF region was replaced via homology-directed repair with a 24-base pair synthetic ‘landing pad’ sequence containing a functional PAM site, flanked by 500 bp homology arms for precise integration. Donor DNA containing the landing pad sequence and flanking homology regions directs the repair process, resulting in seamless replacement of the native gene with the synthetic landing pad.

Landing pad sequences were designed in a previous study^4^ using the Benchling gRNA Design Tool to ensure: (1) presence of a functional NGG PAM motif for future Cas9 targeting, (2) absence of homology to any native yeast genomic sequences, and (3) unique identity for each target gene. This approach provides reversibility, as the resulting edited genome retains the PAM site within the landing pad, enabling future CRISPR targeting for restoration of wild-type gene sequences or further modifications. Targeting efficiency for single gene deletions ranged from 25-75% of colonies showing successful deletion.

### Gene Fragment Design and Cluster Assembly Strategy

Each gene was PCR-amplified from wild-type BY4741 genomic DNA to include its expected upstream and downsteam regulatory elements. However, this posed a challenge as the local genomic environment in yeast is known to play a critical role in transcription regulation in some cases, and moving genes from the DNA context within which they evolved can alter expression even when full-length promoters are included^5^. Furthermore, the ‘neighbouring gene effect’ was also a concern, as deletion or perturbations of genes can disrupt expression of adjacent genes, and altering transcriptional arrangements (tandem, convergent, or divergent orientations) also affects gene expression levels^6^.

The standard design for gene relocation used ∼1000 bp upstream sequences (promoter region) and ∼500 bp downstream sequences (terminator region) to preserve transcriptional regulation during gene relocation (**Figure S1.1B**). These lengths were based on the literature: while most yeast promoters have a median length of 455 bp^7^ with transcription factor binding sites typically enriched ∼115 bp upstream of transcription start sites^8^, the 1 kb upstream region ensures retention of alternative transcription start sites and unknown functional regions. Similarly, the 500 bp terminator region design encompasses yeast 3’ UTRs (median 104-169 bp)^9,10^

Transcriptional gene boundaries were visually validated by using available RNA-Seq expression data shown as tracks on www.yeastgenome.org. In cases where target genes shared intergenic regions with non-target genes (divergent or convergent arrangements), promoter and terminator lengths were adjusted to avoid disrupting neighbouring gene regulation while preserving essential UTR sequences. These gene-specific modifications were aiming to preserve gene expression levels while preventing unintended transcriptional interference.

Synthetic connector sequences (∼150 bp) containing loxP sites were designed for placement between genes to enhance experimental flexibility and enable Cre-mediated deletions. These connectors were extended with gene-specific homology overhangs via PCR to direct *in vivo* assembly order during yeast transformation. The cluster design included a CRISPR landing pad at the 3’ end to enable iterative targeting for subsequent cluster expansions.

### Auxotrophic Markers for Iterative Cluster Expansion

Cluster assembly employed a marker-swapping strategy using three auxotrophic markers (*URA3*, *LEU2*, and *HIS3*) to enable iterative cluster expansion. This method, called SwAP-In (Switching Auxotrophies Progressively by Integration), was originally developed in the Sc2.0 project to replace the native chromosomes with their synthetic versions^11,12^. In our cluster construction, each cluster expansion round integrated new genes while simultaneously replacing the previous selection marker with a different auxotrophic marker, allowing selection for the new cluster version and distinction from previous strains.

This approach leveraged yeast’s proficiency in homologous recombination for *in vivo* assembly of multiple DNA fragments. The assembly process involved co-transforming individual DNA parts (gene fragments, connector sequences, and auxotrophic markers) as separate linear molecules, each engineered with homology ends (∼50 bp) that directed their assembly order. During transformation, yeast cells utilise their endogenous homologous recombination machinery to recognise these overlapping sequences and assemble the fragments into a continuous cluster construct at the target chromosomal locus. This eliminates the need for *in vitro* ligation steps and allows for simultaneous integration and assembly in a single transformation event. The efficiency of this process depends on the length of homology regions, with longer overhangs generally providing more reliable assembly. When this multi-part assembly approach occasionally failed (as observed during 6-gene cluster construction), pre-assembly of the cluster parts into a plasmid followed by linearisation provided an alternative strategy for successful cluster expansion.

The complete cluster construction strategy demonstrates the stepwise assembly approach, progressing from 3-gene to 9-gene clusters through iterative marker swapping. Each expansion round maintained the same genes within the cluster while incorporating additional cell cycle genes. The progression shows the successful integration of *CLN2*, *CLB2*, and *FKH2* in the initial 3-gene cluster, followed by addition of *CLB3* and *CLB5* (5-gene), *SIC1* (6-gene), and finally *CLB4*, *CLN1*, and *FKH1* (9-gene cluster). Each cluster version utilised different auxotrophic markers (*URA3*, *LEU2*, *HIS3*) to enable selection and distinguish successive iterations.

### Construction of the 3-Gene Cluster

The initial 3-gene cluster was established following deletions of the first three genes (*CLN2*, *CLB2*, and *FKH2*), which were individually deleted from their native chromosomal locations using CRISPR/Cas9 and replacing them with the landing pad sequences. This deletion order was chosen based on the published data from synthetic genetic array analysis^13,14^ supporting our hypothesis of the minimal negative interactions between these genes.

Using one of the yeast single knockout strains we had created, *Δclb2*, we next moved to simultaneous deletion of two more genes (*CLN2* and *FKH2*), by supplying four guide RNAs at a time. This resulted in four colonies growing on the selective media, with one later confirmed as a correct triple deletion. Following successful deletion, the three genes were assembled into a synthetic cluster at the *URA3* locus on chromosome V. Gene fragments, connector sequences, and the *URA3* marker were co-transformed as separate parts with designed homology overhangs, enabling one-step *in vivo* assembly via homologous recombination. The initial assembly contained a mutation in a loxP site that was detected by Sanger sequencing which we subsequently corrected by cluster reassembly with a new replacement connector, resulting in the final 3-gene cluster strain.

### Expansion to the 5-Gene Cluster

The 3-gene cluster was expanded through deletion and integration of *CLB3* and *CLB5*. Initial attempts at simultaneous double deletion failed, necessitating sequential individual deletions. *CLB5* was deleted first, followed by *CLB3* deletion in the resulting strain.

For cluster integration, *CLB5* was cloned with a 400 bp terminator and 582 bp promoter to avoid convergent overlap with neighbouring genes and exclusion of a cysteine tRNA gene. *CLB3* was cloned with a 225 bp terminator due to the short intergenic region with the neighbouring *MSH5* gene. The expansion was achieved by CRISPR-mediated targeting downstream of the landing pad in the 3-gene cluster, with homologous recombination incorporating the two new genes and swapping the *URA3* marker to *LEU2*. The resulting 5-gene cluster spanned approximately 18 kb.

### Construction of the 6-Gene Cluster

The 6-gene cluster was constructed by deleting *SIC1* and integrating it into the existing 5-gene cluster. Initial attempts to delete *SIC1* using standard CRISPR approaches with sgRNAs targeting the 5’ and 3’ ends of the ORF failed, likely due to disruption of the neighbouring essential *BOS1* gene located <300 bp away. An alternative sgRNA targeting closer to the start codon also failed to produce viable deletions.

We then used sgRNA to target the start codon region and introduce a premature stop codon rather than complete ORF deletion. A 100 bp synthetic donor DNA was designed to replace the ATG start codon with a TAG amber stop codon and introduce a single nucleotide insertion, creating both a premature stop and frameshift mutation to eliminate functional Sic1 protein production. This approach successfully generated Δ*sic1* cells, though they exhibited slow growth and small colony phenotypes.

For cluster integration, initial attempts using separate gel-purified parts (ConL4, *SIC1* gene fragment, ConL10, and *URA3* marker) with homology overhangs failed to yield any colonies. To troubleshoot this, a *SIC1* ectopic expression plasmid was successfully transformed, demonstrating that Sic1 expression was not problematic. The issue was resolved by pre-assembling the components *in vitro* using Gibson assembly to create a single 3581 bp fragment (ConL4-SIC1-ConL10-HIS3) flanked by NotI sites. This pre-assembled construct was linearised and successfully integrated into the 5-gene cluster, swapping the marker from *LEU2* to *HIS3* and creating a 6-gene cluster of ∼20 kb total length. Whole genome sequencing confirmed cluster integration and revealed a single nucleotide polymorphism in the *CLN2* 5’ UTR region.

### Construction of the 9-Gene Cluster

The final successful cluster expansion created a 9-gene cluster through deletion and integration of *CLB4*, *CLN1*, and *FKH1*. This required individual gene knockouts performed in three rounds, as multiple simultaneous deletions had proven difficult in previous steps.

Round 1 created three strains with seven gene knockouts and six knock-ins (7KO6KI), each containing a single additional deletion: strain yAM021 (Δ*clb4*), strain yAM022 (Δ*fkh1*), and strain yAM023 (Δ*cln1*). Round 2 involved six transformations testing pairwise deletions: deletion of *FKH1* and *CLN1* in the Δ*clb4* strain, deletion of *CLB4* and *CLN1* in the Δ*fkh1* strain, and deletion of *CLB4* and *FKH1* in the Δ*cln1* strain.

This generated the following 8KO6KI strains: Δ*clb4* + Δ*cln1* (yAM025), and Δ*clb4* + Δ*fkh1* (yAM024). However, we had been unable to delete *CLB4* in the *Δcln1* and *Δfkh1* background for unknown reasons, and it would be interesting to investigate this further to confirm whether the order of gene deletions matters.

Round 3 successfully achieved triple deletion: we used the strains yAM024 (*Δclb4* + *Δfkh1*) and yAM025 (*Δclb4* + *Δcln1*) to do single knockouts deleting either *CLN1* or *FKH1*, respectively. Both transformations were successful, yielding the final 9KO6KI strain (yAM026) with triple deletion (Δ*clb4*, Δ*fkh1*, Δ*cln1*).

For cluster integration, gene fragments required specific modifications based on their genomic contexts: *FKH1* was cloned with only a 250 bp core promoter due to the shared 547 bp intergenic region with the neighbouring transcriptional regulator *ASG1*, containing divergent promoters for both genes. The predicted 5’ UTR boundary for *FKH1* is 146 bp, ensuring the core regulatory elements were captured. *CLB4* required both shortened promoter (250 bp) and terminator (350 bp) sequences due to its tight genomic clustering with *PNP1* (276 bp upstream) and *ATG38* (402 bp downstream), with predicted core UTR regions of only 58 bp (5’) and 174 bp (3’).

The three genes were integrated using the same assembly strategy used for previous cluster expansion rounds. A double-strand break was introduced after connector ConL10 using CRISPR/Cas9, with donor DNA supplied as separate gel-purified gene fragments and connectors bearing PCR-added homology overhangs. The transformation successfully yielded colonies assembling the complete 9-gene cluster totalling approximately 28 kb. Four colonies positive for both the CRISPR plasmid marker (*URA3*) and the new cluster auxotrophic marker (*LEU2*) were screened, with three showing correct assembly by colony PCR verification of gene-connector junctions. PCR products of expected sizes were confirmed by gel electrophoresis and validated by Sanger sequencing, demonstrating accurate assembly of all transformed components. The final strain (yAM027, designated 9KO9KI) was sequenced using Illumina genome sequencing.

### Challenges to Expand Beyond 9 Genes

Multiple approaches to delete and integrate *CLB1* and *CLB6* failed due to their semi-essential nature and due to technical complications. Attempts at simultaneous deletion of both resulted in unfaithful recombination events with partial *CLB1* ORF deletions while leaving *CLB6* intact. Individual deletion attempts using various sgRNA combinations yielded only wild-type colonies.

Alternative approaches including integration before deletion also failed. Assembly of *CLB1* and *CLB6* gene fragments into the cluster was unsuccessful, with nanopore sequencing revealing incorrect integrations with missing gene segments. This suggests homologous recombination issues in the 9-gene cluster strain.

As an alternative deletion strategy, KanMX4 cassette disruptive integration into the target CDS was attempted. While *CLB1* deletion was achieved using donor DNA amplified from Euroscarf deletion collection strain for ΔCLB1^15^. *CLB6* deletion failed due to multiple PCR products arising from the DNA of the corresponding ΔCLB6 Euroscarf strain, attributed to chromosomal rearrangements involving the Ty-3 retrotransposon downstream of *CLB6*.

To complete the cluster, we also suggest integrating the *CLN3* gene, in addition to the *CLB1* and *CLB6* genes we were working on. The vision is therefore for a future 12-gene cell cycle cluster that encompasses a full set of genes responsible for encoding the ‘waves of cyclins’ function.

### Verification of CRISPR/Cas9-mediated construction of synthetic cell cycle gene clusters

Each construction step was verified using complementary approaches ensuring genomic accuracy:

#### Colony PCR screening

Locus-specific primers targeting integration junctions and deletion sites provided initial confirmation of successful modifications.

#### Sanger sequencing

Sanger sequencing: Genomic DNA spanning ∼100 bp upstream and downstream of each edited region, including the homology arms of the donor DNA, was PCR-amplified with Phusion or Q5 high-fidelity DNA polymerase. The resulting amplicons were Sanger sequenced (Source Bioscience) using the amplification primers, with additional internal primers added where the edited region exceeded the read length, in order to obtain full coverage of the edited locus. Reads were aligned to the expected sequence on Benchling, verifying precise junction sequences at gene knockouts and at gene–connector boundaries within the assembled cluster, and confirming the absence of mutations at editing sites.

#### Whole genome sequencing

Whole genome sequencing: Illumina paired-end sequencing was performed by SeqCentre (formerly MIGS, Pittsburgh, PA, USA) on an Illumina platform. Raw reads were quality-controlled and adapter-trimmed with fastp, assessing base quality scores, GC content, and read length to flag low-quality bases and contaminants. Pre-processed reads were aligned with BWA-MEM/BWA-MEM2 to a haploid BY4741 reference genome modified to incorporate our designed CRISPR edits, with SAM-to-BAM conversion, indexing and statistics performed via Samtools and the Galaxy platform. Alignments were visualised in the Integrative Genomics Viewer (IGV) and per-chromosome coverage was plotted using the BAM Coverage Plotter tool. This workflow allowed us to confirm correct cluster integration at the chromosome V URA3 locus, verify uniform coverage at each landing pad, identify any local mutations, SNPs or coverage drops, and screen for off-target effects across the rest of the genome.

### Whole Genome Sequencing Analysis

#### 6-Gene Cluster Validation

Illumina whole genome sequencing of strain yAM018 (6KO6KI) was performed using a modified BY4741 reference genome incorporating designed CRISPR edits. Quality control using fastp showed 91% of reads were error-free with phred scores >30, with variable base content in the first 20 bases suggesting minor adapter contamination. Trimmed reads aligned to the reference genome revealed uniform coverage across the 6-gene cluster region at the chromosome V *URA3* locus, confirming correct integration location and gene order.

A notable coverage drop was observed spanning ∼450 bp from the loxP site in connector ConL5 to ∼300 bp from the 5’ end of *CLB3*, initially raising concerns about potential gaps in cluster construction. However, subsequent targeted Oxford Nanopore Technologies sequencing revealed no gaps, suggesting this was an artifact of short-read alignment. Additionally, a single nucleotide polymorphism was identified in the 5’ UTR of the *CLN2* gene within the cluster, later determined to be cryptic as it was absent in the 9-gene cluster strain derived from this parent.

Comprehensive analysis of all six gene knockout loci demonstrated uniform coverage and confirmed precise editing according to design, with no local mutations or off-target effects detected. Each landing pad integration was verified at the targeted genomic regions for *CLN2*, *CLB2*, *FKH2*, *CLB3*, *CLB5*, and *SIC1*.

#### 9-Gene Cluster Validation

Illumina sequencing of strain yAM027 (9KO9KI) showed 95.5% of reads achieving phred scores >30. WGS confirmed successful integration of the complete 9-gene cluster (∼28 kb) at chromosome V with correct gene order and loxP-flanked connector sequences, however, the previously observed coverage drop persisted in the same location.

Validation of all nine gene knockout loci revealed uniform coverage and precise editing for eight of nine targets. The exception was the *FKH1* locus, where an 8-base pair gap was detected at the 3’ end of the landing pad sequence, with these bases inserted into the 5’ end of the spacer and loss of the PAM site (NGG). While this mutation does not compromise current functionality, it could complicate future reversion attempts, though new sgRNAs could be designed to target the altered site. The absence of the *CLN2* SNP in the 9-gene cluster confirmed its cryptic nature in the parent strain.

### Conclusions

This work demonstrates the successful application of CRISPR-based genome engineering for ‘defragmenting’ the yeast genome by relocating functionally related genes into synthetic clusters. The successful assembly of a 9-gene cell cycle cluster (∼28 kb) represents a significant achievement in eukaryotic genome engineering. loxP sites in the cluster enable controlled gene deletions *in vivo* using an estradiol-induced recombinase enzyme, Cre-EBD, providing a valuable platform for studying minimal gene networks.

## Notes

### Competing Interest Statement

The authors have declared no competing interest.

